# Context-dependent peptide recognition shapes tyrosine kinase substrate specificity beyond consensus motifs

**DOI:** 10.64898/2026.05.10.724103

**Authors:** Hannah Edstrom Athol, Annette Thompson, Nolan O’Connor, Michael R. Shirts, Joel M. Kralj, Jerome M. Fox

## Abstract

Protein tyrosine kinases (PTKs) regulate cellular biochemistry by phosphorylating tyrosine residues that alter protein function; their substrate preferences define the topology of signaling cascades. Previous studies of PTKs have mapped their average preferences for amino acids surrounding phosphorylation sites, but their sensitivity to sequence variation remains poorly understood. Here, we used microbial biosensors for PTK activity to examine the influence of local sequence context on substrate specificity. Across five well-studied PTKs, we identified amino acid substitutions within consensus substrates that could confer sensitivity to substrate length or enhance selectivity for one PTK over others. Using a secondary decoy screen, we found sequence-diverse substrates with unexpectedly orthogonal PTK compatibilities. Our findings show how context-specific sequence features alter PTK substrate specificity far beyond what might be expected from classical consensus models and establish an experimental framework for defining the limits of substrate overlap between closely related kinases.

## INTRODUCTION

Protein tyrosine kinases (PTKs) catalyze the ATP-dependent phosphorylation of tyrosine residues and are centrally important to the survival of eukaryotic cells.^1^ They regulate numerous biological processes (e.g., cellular metabolism,^2,3^ migration,^4^ proliferation,^5^ memory,^6^ and immunity^7–9)^ by phosphorylating tyrosine motifs that alter protein activity, complexation, and localization. Anomalously regulated PTKs cause a broad spectrum of diseases, commonly cancer.^3,10–13^ Today, over 60 approved drugs target these enzymes, and many more are in development.^14^ Despite the importance of PTKs in biology and medicine, the molecular determinants of their regulatory interactions remain only partially understood.

High-throughput biochemical methods have enabled broad profiling of the intrinsic substrate specificities of PTKs. In the most common approach, purified PTKs are assayed on degenerate libraries of short peptides (e.g., 10-15 residues) to produce maps of amino acid preferences at each position.^15–17^ In a recent tour de force, Johnson and colleagues used a version of this approach, termed positional scanning peptide array (PSPA) analysis,^18,19^ to profile the substrate specificity of all human PTKs.^15^ When supplemented with bioinformatic analyses, heatmaps of PTK-specific amino acid preferences allowed them to identify compatible PTKs for select sites from the human Tyr phosphoproteome, including known PTK-substrate pairs. Interestingly, the approach struggled to detect autophosphorylation sites, which are subject to induced proximity (e.g., PTK dimers) and intermolecular conformational constraints.

The advent of low-cost sequencing has enabled the use of genetically encoded peptide libraries to map substrate specificity with extraordinary throughput and precision.^9,20,21^ In one seminal study, Kuriyan and colleagues used bacterial peptide display (BPD) to profile the activity of EGFR kinase on a peptide library containing established human phosphosites.^22^ In this study, autophosphorylation sites in the tail of EGFR did not match expected consensus motifs but, instead, appeared to balance EGFR compatibility and c-Src incompatibility. Follow-up mutational analyses of tail sites showed that preferences at the -1-position depended on the identity of neighboring residues, perhaps because of multiple binding modes. Other studies have observed similar dependencies^23,24^. These results highlight a key limitation of consensus maps: the assumption that amino acid preferences at each position are independent of the sequences in which they occur. A detailed understanding of this context dependence (i.e., the influence of amino acid position, substrate length, and epistatic effects on the site-specific amino acid preferences of a given PTK) is essential for using sequence data to map signaling cascades.

In this study, we used a genetically encoded biosensor for PTK activity to explore the influence of sequence context on substrate specificity. Our biosensor, which links PTK activity to the expression of a gene for antibiotic resistance in *Escherichia coli* (*E. coli*), enabled the construction of position-specific scoring matrices (PSSMs) consistent with prior work and facilitated rapid quantitation of substrate compatibility via simple growth experiments. In general, consensus motifs were most compatible with cognate kinases (relative to the other PTKs) but obscured highly influential sites where single amino acid substitutions could disrupt PTK compatibility or dramatically enhance selectivity for one PTK over another. By adding a second “decoy”-based screen for incompatible substrates, we identified substrate sequences with orthogonal PTK compatibilities that could not be predicted from classical consensus maps. Our findings show how differences in context-specific amino acid preferences between PTKs can alter substrate specificity far beyond what might be expected from classical consensus models.

## RESULTS

### Genetically encoded biosensors enable rapid profiling of PTK substrate specificity

To profile the activity of PTKs on diverse substrate pools, we developed a high-throughput assay that uses a bacterial two-hybrid (B2H) system to detect PTK activity in *E. coli* (Fig. 1A). In this B2H, PTKs phosphorylate a peptide, allowing it to bind to an SRC homology 2 (SH2) domain, and the resulting phosphopeptide-SH2 complex localizes RNA polymerase to a gene for spectinomycin resistance (specR). We have used similar systems to guide inhibitor biosynthesis^25,26^ and to study neutral drift in protein tyrosine phosphatases;^27^ others have used them for mutational analyses of PTKs.^28^ We chose to use a B2H for the present study because it avoids PTK purification, does not require specialized reagents or instrumentation, and enables rapid verification of substrate compatibility via simple growth experiments.

**Figure 1.**
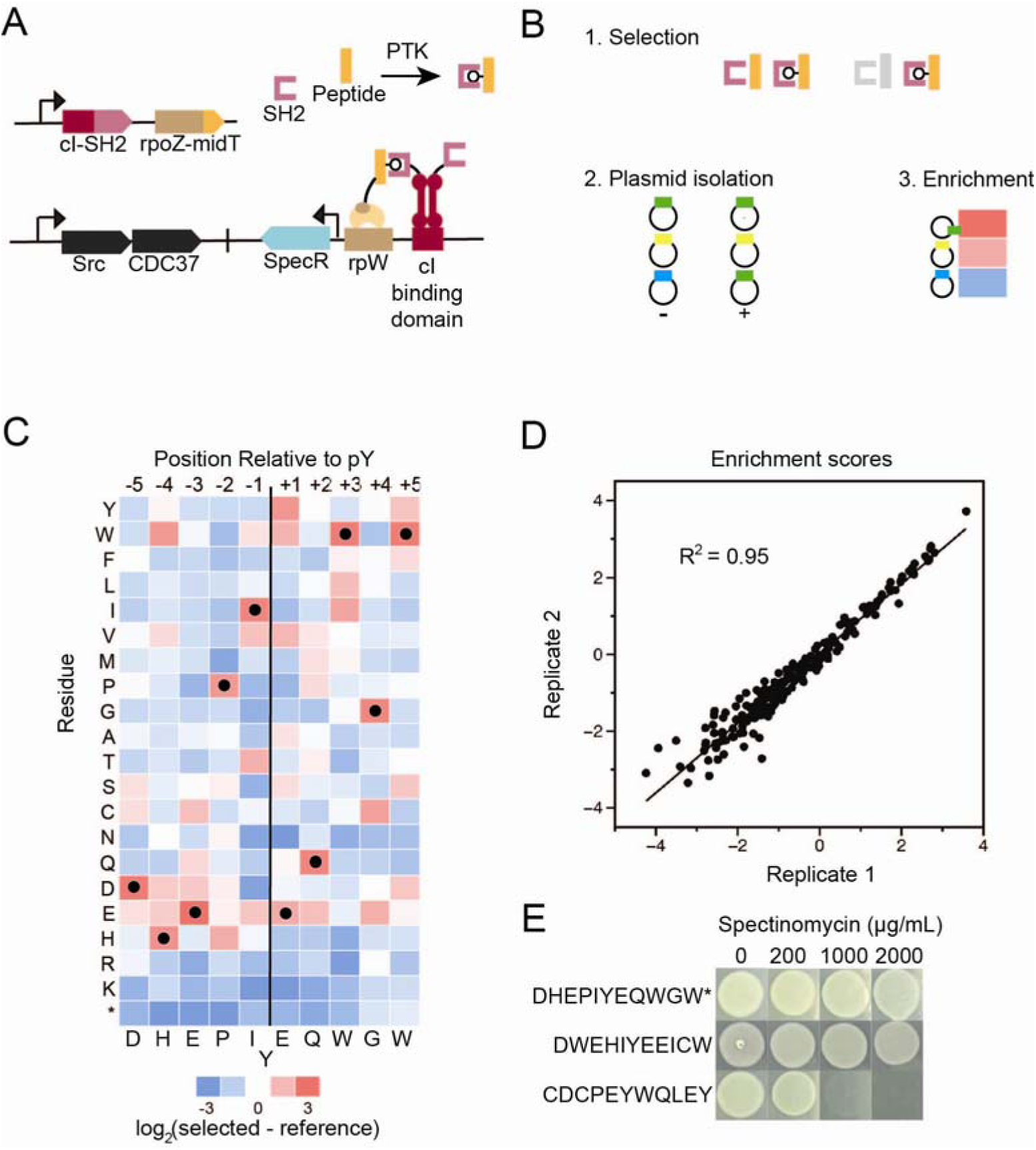
Genetically encoded biosensors enable rapid profiling of substrate specificity. (A) To detect kinase activity, we used a bacterial two-hybrid (B2H) system based on two protein fusions: (i) a peptide substrate fused to the l2 subunit of RNA polymerase (RPl2) and (ii) a “superbinder” SH2 domain fused to a cI repressor. Phosphorylation of the peptide enables peptide-SH2 binding and localizes the RNA polymerase to a gene for spectinomycin resistance (SpecR). (B) We used the B2H to screen substrate libraries (X_5_-Y-X_5_, X = any one of the 20 common amino acids or a stop codon, *) for PTK compatibility, which yielded enrichment on spectinomycin plates. (C) A position-specific scoring matrix (PSSM) for c-Src shows the enrichment of amino acids at each position within an X_5_-Y-X_5_ substrate, accounting for stop codons (mean, n = 2 biological replicates). Closed circles, which mark the most enriched residues at each position, define the consensus motif (Fig. S9, Table S7). (D) Enrichment values were consistent between biological replicates. (E) B2H-based assessment of the compatibility of substrates defined by the (rows 1-2) most enriched or (row 3) second most enriched residues at each position, when stop codons are either (*) included or (no asterisk) excluded from the enrichment calculation. Data represent averages of (C) n = 2 or (E) n = 3 biological replicates.

Our B2H uses an SH2 domain, which could exert its own substrate specificity. To minimize this effect, we undertook three precautions: (i) We used a highly permissive domain—the SH2 of c-Src kinase—and added three “superbinder” mutations known to enhance binding affinity and broaden specificity.^29–31^ This SH2 variant has outperformed anti-pTyr antibodies in phosphoproteomics analyses.^32^ (ii) We focused our analyses on comparisons in which the influence of the SH2 domain is constant between experiments. (iii) We confirmed the lack of significant bias by using our method to check PTK compatibility for substrates identified with alternative methods (Figs. S1 and S2).

We validated our approach by profiling the substrate specificity of c-Src, a well-studied PTK with established substrate profiles^15,29^—a good positive control. Briefly, we used degenerate primers to construct an 11-residue substrate library (X_5_-Y-X_5_, where X is any of the 20 common amino acids or a stop codon) and transformed it into cells harboring the B2H plasmid. We grew cells in the presence and absence of antibiotic, sequenced substrate populations from each condition, and estimated the log_2_-fold enrichment for every amino acid at each position (Figs. 1B-1C). Position-specific enrichment values showed excellent reproducibility between biological replicates (Figs. 1D, S1, S2) and good agreement with previous PSSMs (Fig. S5).

**Figure 2.**
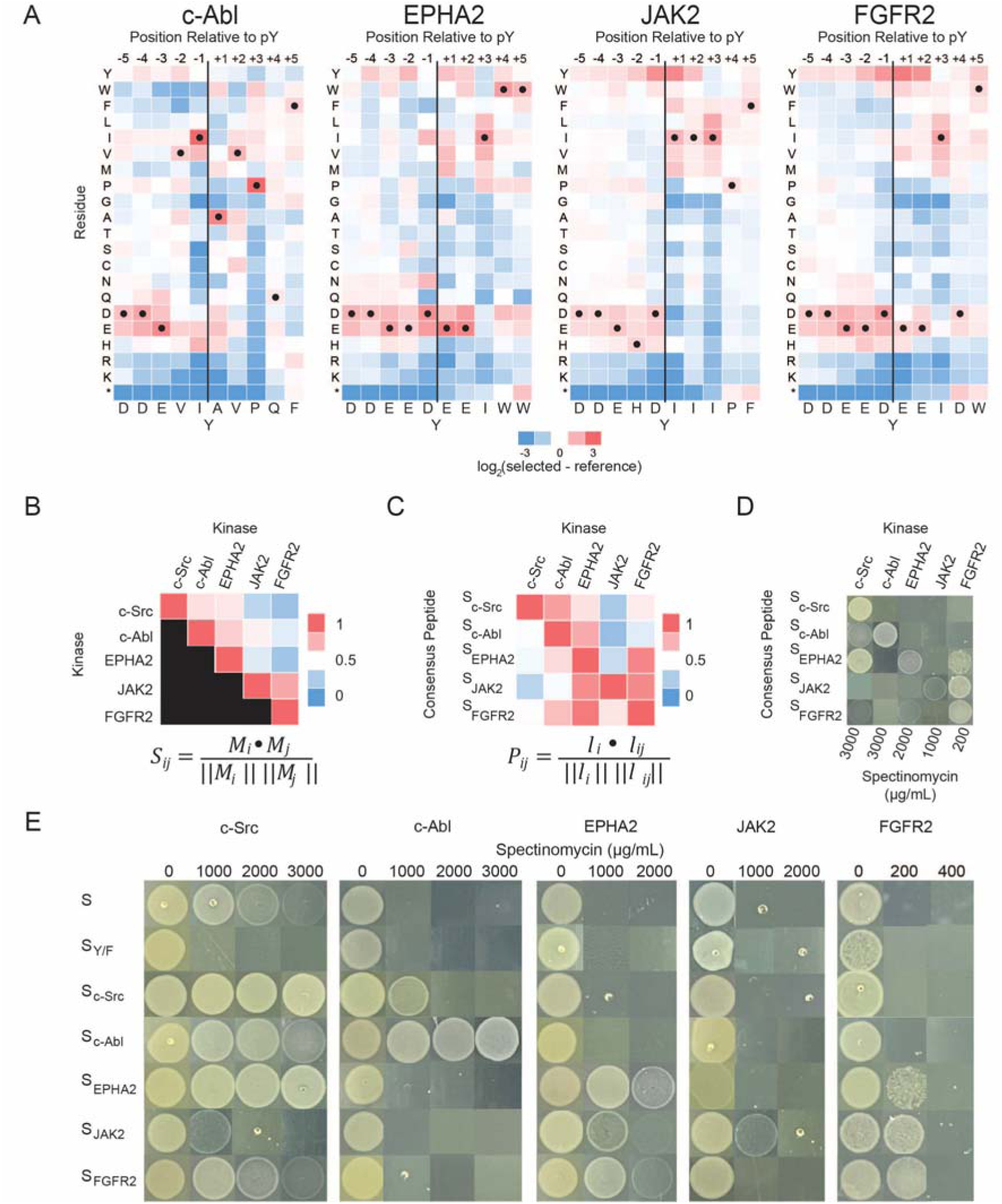
Heatmaps reveal critical differences in substrate scope. (A) PSSMs for additional PTKs. (B) A comparison of global substrate preferences of each binary combination of PTK. *S_ij_* is the cosine similarity of *M_i_* and *M_j_*, the PSSMs of PTKs *i* and *j*. (C) A comparison of substrate compatibility between PTKs. *P_ij_* is the cosine similarity of *l_i_*, a vector of PTK*i* enrichment values for the PTK*i* consensus, and *l_ij_*, a vector of PTK*j* enrichment values for the same sequence. (D) B2H-based assessment of consensus motif compatibility with each PTK, based on the maximum resistance enabled by cognate motifs for each PTK. (E) B2H-based assessment of consensus motif compatibility with each PTK. Controls: S and S_Y/F_ denote positive and negative controls for c-Src. In A, data represent averages of n = 2 biological replicates (Fig. S6). In D and E, images depict drops from n = 3 biological replicates (Fig. S7).

As with most other PTKs, c-Src favored N-terminal acidic residues (e.g., aspartate at −5 and glutamate at −3), relative to the phosphoacceptor Tyr, and disfavored basic residues at all positions. The preference for N-terminal acidic residues complements a positively charged region of the active site—a likely interaction point, based on alignments with other PTK complexes (Fig. S6; ^33–37)^. As reported in prior work, c-Src showed a strong preference for isoleucine at −1 and hydrophobic residues at +3; we also observed previously established—though more moderate—preferences at −2 and +2 (i.e., proline and glutamate)^24,38^. Positive enrichment for histidine at −2 is surprising, based on prior data, and could reflect the influence of c-Src SH2, which favors this substitution;^29,30,39^ however, superbinder mutations dramatically weaken this specificity.^32,40,41^ In general, the consensus preferences measured by our approach are similarly consistent with alternative methods as those methods are with each other (Fig. S5), suggesting that PTK-specific substrate preferences are the dominant contributor to enrichment profiles.

We used our B2H system to confirm the compatibility of motifs suggested by our PSSM. To begin, we examined two consensus motifs defined by the most enriched residue at each position, as calculated by either (i) including or (ii) excluding stop codons in enrichment estimates; these two motifs had similar sequences and conferred similar levels of spectinomycin resistance with our B2H system (Fig. 1E, S7, S8). Next, we tested a substrate defined by the second most enriched residue at each position. This substrate dramatically reduced resistance, indicating that the extent of amino acid enrichment at each position in a PSSM helps predict the overall extent of substrate compatibility. Moving forward, we defined all enrichment scores by accounting for stop codons in our enrichment estimates.

### PTKs show prominent differences in substrate scope

We explored PTK-specific substrate preferences by examining the kinase domains of c-Abl, EPHA2, JAK2, and FGFR2. These influential regulatory enzymes are sequence diverse (∼25-70% identity in PTK domains) and fall into different specificity groups, based on prior use of hierarchical clustering to group human PTKs by substrate specificity (Table S1)^38,42^. In our system, these PTKs yielded different levels of maximal antibiotic resistance (i.e., different selection conditions)—a discrepancy that we attributed to differences in PTK expression and associated toxicity when expression was too high (Fig. S9). Nonetheless, all were compatible with our screen; as with c-Src, the PSSMs were highly concordant with those based on alternative approaches (Figs. 2A, S5, and S10), where previously determined consensus motifs showed predictable compatibilities in our assays (Fig. S2). In general, all four enzymes favored N-terminal acidic residues, disfavored basic residues throughout, and, except for JAK2, preferred isoleucine at −1; these specificity patterns are consistent with prior reports.^15,29^ Notably, preferences for histidine at −2 2 varied across PTKs, indicating that enrichment at this position, even if influenced by the SH2 domain, was constrained by each PTK.

Having validated the use of our B2H system to profile PTK substrate specificity, we explored differences in substrate preferences. First, we calculated the cosine similarity (*S_ij_*) of enrichment patterns for each binary combination of PTKs (Fig. S11). In Eq. 1, *M_i_* and *M_j_* are the

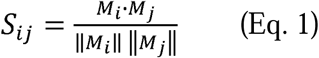

enrichment maps for PTKs *i* and *j*, and *S_ij_* is a metric for the similarity of those maps. This metric weighs all positions equally and assumes symmetric preferences between PTKs; that is, a value of *S_ij_* close to 1 suggests that substrates are similarly compatible with PTKs *i* and *j*. Values of *S_ij_* suggested two specificity groups: (i) c-Src, c-Abl, and EPHA2, and (ii) JAK2 and FGFR2 (Fig. 2B). We used our B2H system to test the predictive power of *S_ij_* on consensus motifs (Figs. 2E and S12). In general, consensus motifs were highly compatible with their cognate kinases but showed inconsistent compatibilities within specificity groups. For example, the consensus motifs for c-Src, c-Abl, and EPHA2 conferred similar antibiotic resistance for c-Src but not EPHA2, which preferred its own motif. JAK2 also showed a strong preference for its consensus, even though FGFR2 worked similarly well with FGFR2, JAK2, and EPHA2 motifs. The asymmetric substrate compatibility between PTKs highlights differences in their promiscuity (i.e., the extent of substrate preference) that are not obvious from simple comparisons of their specificity maps.

Next, we sought a metric to assess differences in PTK compatibilities with a specified consensus sequence. In Eq. 2, *l_i_* is a vector of the enrichment values for each amino acid in the

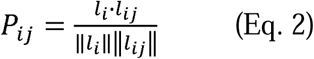

consensus motif for PTK_i_; *l_ij_* is a vector of enrichment values for the same sequence for PTK_j_; and *P_ij_* describes the compatibility of the PTK_i_ consensus motif with PTK_j_ (Fig. 2C). When compared to *S_ij_*, this metric was more predictive of the narrow and broad substrate compatibilities of JAK2 and FGFR2; however, it did not predict the promiscuity of c-Src (Figs. 2C and 2D)—a shortcoming shared by previously explored metrics for substrate compatibility (Table S2). These findings suggest that specific numerical enrichment values can have very different compatibility implications for the PSSMs of PTKs with different substrate scopes.

### PTK-substrate interactions show strong positional dependence

Differences in the range of acceptable amino acids (or stop codons) at each substrate site indicate that some sites are more influential than others (Fig. 3A). To explore this positional dependence, we averaged the enrichment of various side chain functionalities at each site (Fig. 3B and S13). PTKs showed prominent differences in their preferences for aromatic and acidic residues at the −1 site and C-terminus, an indication that these regions have a disproportionate influence on substrate specificity (Fig. S14). Indeed, when we swapped the N- and C-termini of consensus motifs for c-Src and c-Abl, PTK compatibility followed the C-terminal sequence (Figs. 3C and S15A). To follow up, we prepared chimeric substrates with a “general” acidic N-terminus appended to each C-terminal consensus motif. Once again, PTK compatibility followed the C-terminus—though the chimeric substrates reduced overall antibiotic resistance, perhaps a result of reduced binding affinity (Figs. 3C and S15A). These growth experiments show that differences in the C-terminal sequence alone are sufficient to select for specific PTKs.

**Figure 3.**
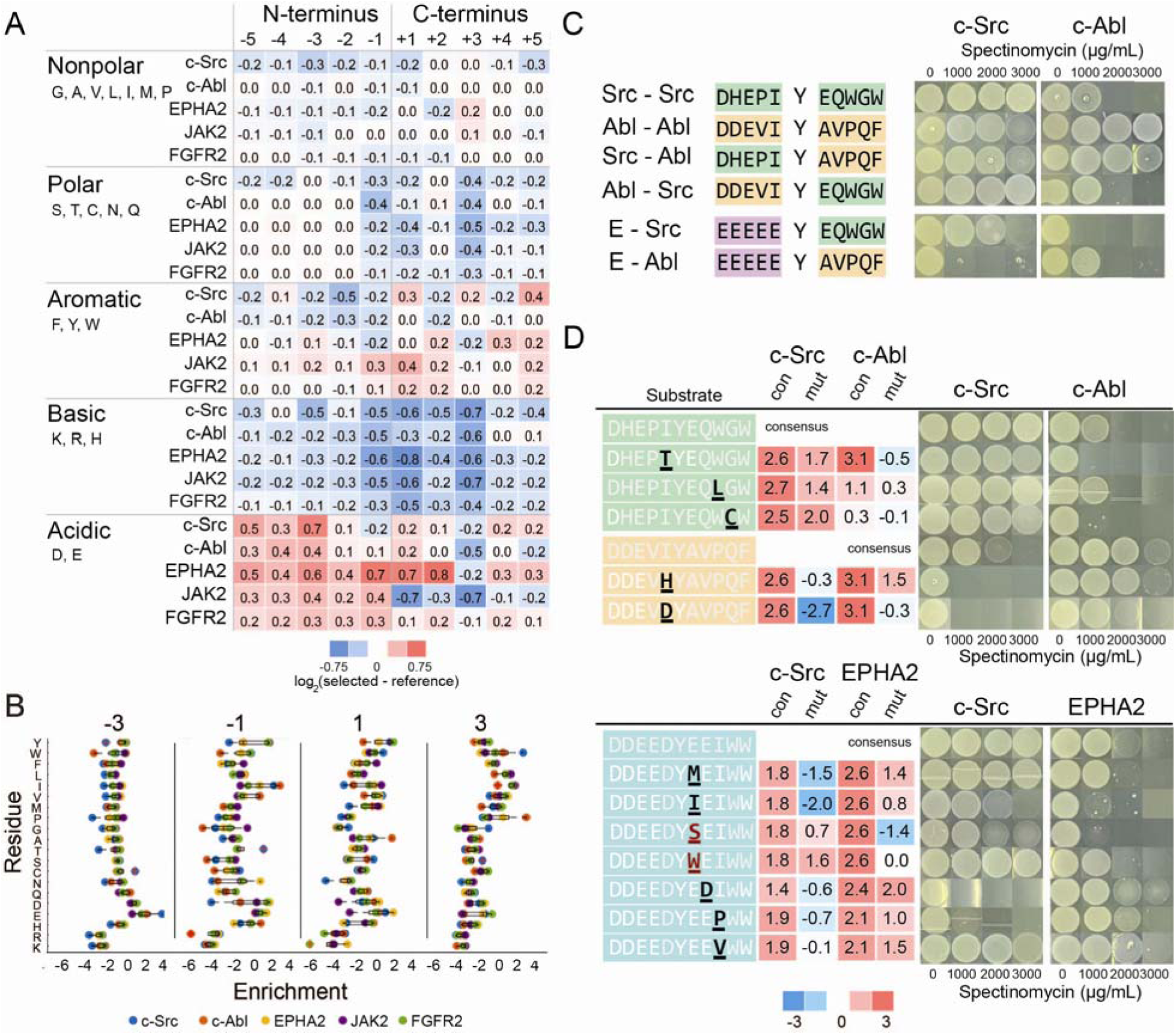
Amino acid preferences show strong positional dependence. (A) The average enrichment of different side chain functionalities at each substrate site (X_5_-Y-X_5_). PTKs showed major differences in their preferences for aromatic and acidic residues at the −1 site and C-terminus. (B) The enrichment of each amino acid at each substrate site (X_5_-Y-X_5_). Preferences differ most near the phospho-acceptor tyrosine. (C) Top: B2H-based assessment of substrate compatibility for substrates constructed with chimeras of the c-Src (green) and c-Abl (gold) consensus motifs suggests that the C-terminus controls PTK compatibility. Bottom: Replacing the N-terminus with “EEEEE” yields similar compatibility differences but reduces overall survival. (D) B2H-based assessment of substrate compatibility for consensus substrates with amino acid substitutions that showed differential enrichment between PSSMs. Numbers denote the log_2_-fold enrichment of the underlined position for c-Src, c-Abl, or EPHA2 as follows: (i) consensus amino acid (con) and (ii) mutation (mut). Mutations across a range of positions (i.e., - 1 to +4) can alter inter-PTK orthogonality, but their impacts are difficult to predict from enrichment values. Data depict (A-B) the averages from n = 2 biological replicates or (C-D) drops from n = 3 biological replicates (Fig. S9).

### Positional preferences are sequence-dependent

To examine the impact of individual mutations on PTK compatibility, we modified consensus motifs with amino acid substitutions that showed differential enrichment between PSSMs (Figs. 3D and S15B). Modifying the consensus motif for c-Src with mutations enriched for c-Src but not c-Abl reduced c-Abl compatibility in two cases (-1Thr and +4Cys) but not a third (+3Leu). With similar consistency, modifying the c-Abl motif with -1His, which was enriched for c-Abl, reduced c-Src compatibility, but adding -1Asp, which was depleted for c-Src, reduced compatibility with both PTKs. Remarkably, -1Asp appears in the EPHA2 motif, which is highly compatible with c-Src, suggesting that its impact depends on sequence. Turning our attention to this motif, adding amino acids enriched for EPHA2 but depleted or neutral for c-Src brought about either (i) a large, disproportionate reduction in c-Src compatibility (+2Asp and +3Pro), (ii) no impact on PTK compatibility (+1Met and +3Val), or (iii) reduced compatibility with both PTKs (+1Ile). Similarly, mutations enriched for c-Src but not EPHA2 reduced compatibility with both PTKs (+1Ser) or had no effect (+1Trp). Overall, our B2H studies show that individual mutations across a range of sites (i.e., −1 to +4) can impact PTK compatibility but indicate that the extent of impact is difficult to predict from differences in enrichment values alone.

Deep mutational scanning (DMS) is a powerful method for examining the influence of mutations on protein structure and function.^43,44^ We probed sequence context more directly by carrying out DMS of the consensus motifs of c-Src and JAK2 (Figs. 4A, S16). These PTKs had discrepant substrate specificities, and we speculated that they might exhibit distinct sensitivities to consensus modifications. Indeed, c-Src retained its strong preference for -1Ile but favored many substitutions at the −5, −4, +2, and +5 sites, while JAK2 retained its preference for +3Ile but favored substitutions at −1, +1, and +2 sites. The high activity of c-Src on its consensus substrate made changes in compatibility challenging to detect (Figs. 4B, S17, and S18), so we focused on JAK2. Depleted substitutions reduced substrate compatibility, and enriched substitutions preserved or enhanced it. Our DMS analysis illustrates how standard consensus maps can obscure both the strength of PTK preferences at key positions (e.g., the strict preferences of c-Src and JAK2 for −1 and +3) and the breadth of tolerated residues, which can change with sequence.

**Figure 4.**
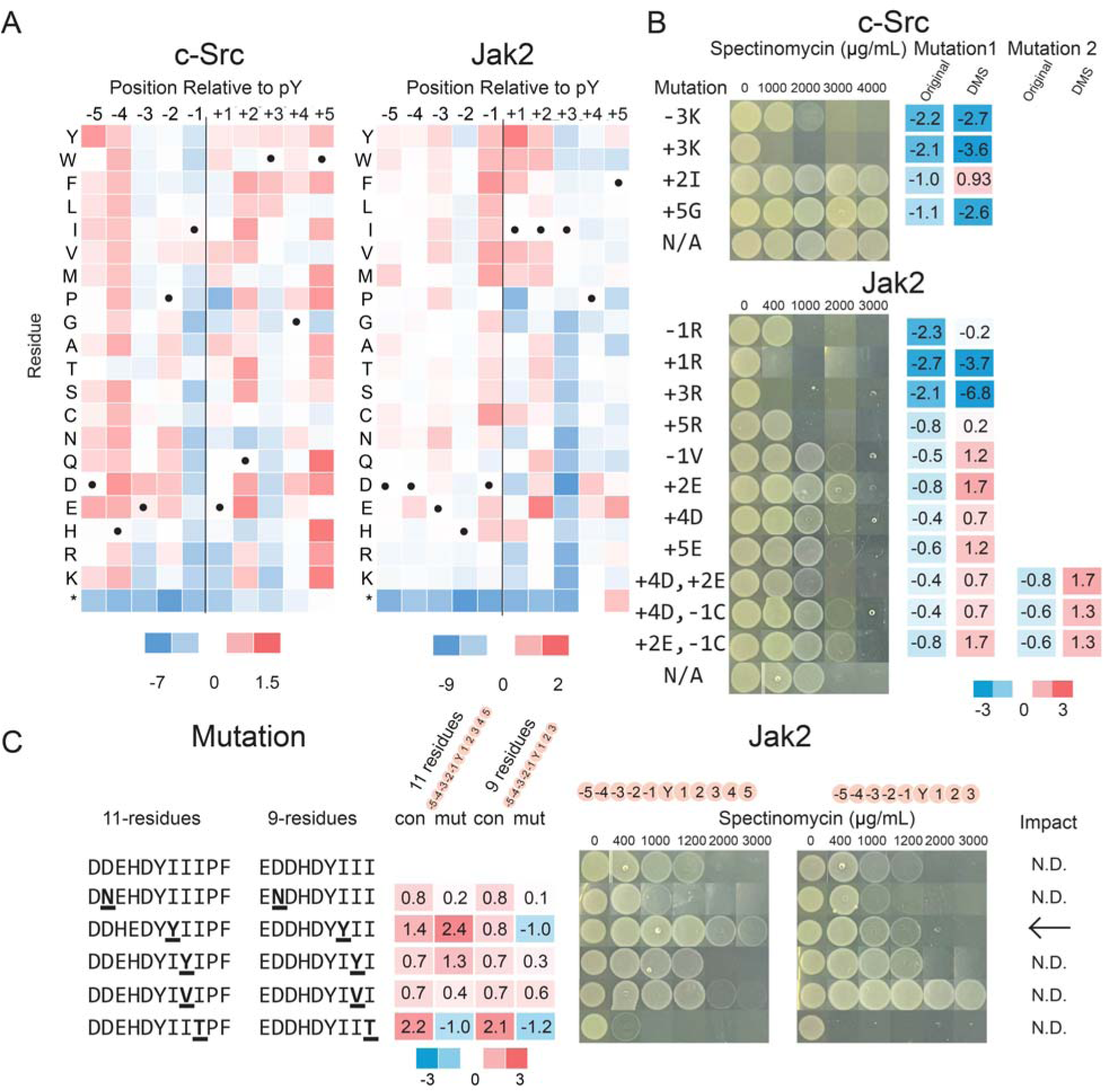
Amino acid preferences depend on sequence context. (A) DMS analysis of consensus motifs for c-Src and JAK2 (black dots denote consensus). Data depict the average of n = 2 biological replicates (Fig. S13). (B) B2H-based assessment of substrate compatibility for consensus motifs with individual mutations, where E*_i_* is the enrichment score for the residue in the original PSSM (Fig. 2A), and E*_f_* is the enrichment score for the residue in the DMS PSSM (Fig. 5B). (C) B2H-based assessment of JAK2 compatibility with its 11-residue consensus motif, a 9-residue truncation of that motif, and single-site mutants. I1Y, a substitution enriched only in the 11-residue library, enhanced JAK2 activity on the longer substrate but not the shorter. N.D. indicates no data. Data depict averages from (A-C) biological replicates, and (B-C) images depict drops from n = 3 biological replicates (Figs. S14 and S15).

### PTKs show different sensitivities to substrate length

JAK2 and FGFR2 were unusually tolerant of stop codons at positions +4 and +5, an indication that these positions are not required for substrate binding. Their activity on varied substrate lengths is consistent with tyrosine enrichment at several sites (-2 through +2), which can effectively shift the location of the phosphorylation site. Multi-site tyrosine enrichment, which was observed in peptide arrays, is also suggestive of multi-site phosphorylation—or “phosphopriming”—a widespread phenomenon in PTKs.^15^ To probe the impact of substrate length more directly, we screened JAK2 and c-Src against 9-residue peptide libraries with the final two sites removed (X_5_-Y-X_3_; Fig. S19). Enrichment was similar between the two libraries for JAK2, but not c-Src, where shorter peptides showed limited compatibility.

We followed up by examining a small subset of JAK2 substrates with drop-based plating. The consensus sequences for the 11- and 9-residue libraries—hereafter S_11_ and S_9_–were similarly compatible with JAK2, and mutations had a range of effects (Figs. 4C, S20, and S1E): +1Tyr, which was enriched for S_11_ and depleted for S_9_, improved growth for only the long substrate; +3Thr, which was consistently depleted, rendered both substrates incompatible; and a handful of mutations with similar or marginally discrepant enrichments between S_11_ and S_9_ had modest effects on growth. Our results indicate that JAK2 prefers YY motifs on substrates with +3Ile but tolerates sequence variation or truncations thereafter (i.e., +4 to +5) and, more broadly, highlight the potential for substrate length to alter the impact of mutations on substrate compatibility.

### In vitro validation of substrate preferences

Our assertion that antibiotic resistance afforded by our B2H systems reports on PTK-substrate compatibility is supported by (i) the consistency between our PSSMs and those generated via previous methods (Fig. S5) and (ii) the compatibility of previously determined consensus substrates with our B2H system (Fig. S2). In preparation for more detailed biophysical studies, we tested this assertion more directly by carrying out two sets of analyses: First, we used drop plating to evaluate PTK-substrate combinations previously examined with in vitro kinetic assays; indeed, trends in antibiotic resistance mirrored those observed in kinetic data (Fig S1). Second, we measured complete Michaelis-Menten curves for six substrates that conferred different levels of antibiotic resistance in B2H systems for c-Src, c-Abl, and JAK2 (Figs. 5A, S1B, and S1C). In general, large differences in antibiotic resistance between closely related substrates (e.g., ∼2000 µg/ml between the maximum tolerated concentrations) translated to significant differences in K_M_: (i) Both c-Abl and c-Src were active on the c-Abl consensus peptide, but only c-Abl had measurable activity on its -1His mutant, as predicted in Fig. 3D. (ii) The K_M_ of JAK2 was higher for the +1Tyr mutant of the S_11_ consensus than for the same mutant of the S_9_ consensus, as in Fig. 4C. (iii) JAK2 activity was similar on substrate pairs where differences in antibiotic resistance were more marginal, notably the consensus and +2Glu mutant discussed in Fig. 4B. These results support the use of drop plating to assess major differences in substrate compatibility between PTKs prior to more detailed biophysical studies (39, 40).

**Figure 5.**
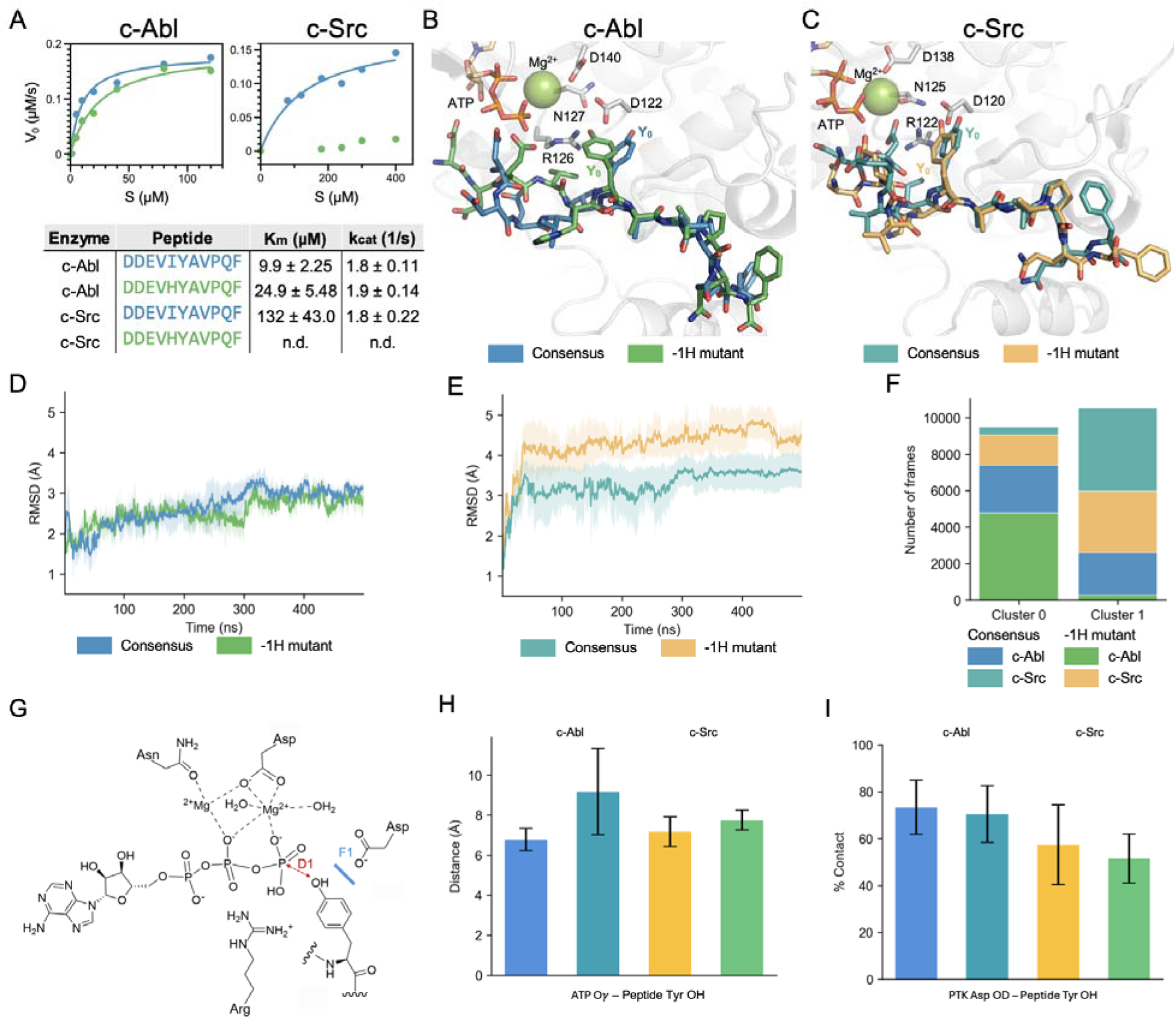
Molecular modeling tools struggle to predict the impact of individual mutations on PTK-substrate compatibility. (A) In vitro kinetic assays of PTK activity on DDEVIYAVPQF (the c-Abl consensus) and DDEVHYAVPQF (the -1His mutant). Consistent with Fig. 3D, c-Abl is compatible with both peptides, but c-Src is only compatible with the consensus peptide. (B-C) Centroid structures of the consensus and the -1His mutant. (D-E) RMSD of peptide backbone atoms from their starting structures for (D) c-Abl and (E) c-Src. Shading is the standard error of the mean for n=3 (Fig. S18). (F) Clustered trajectories of peptide-PTK pairs separated the majority of c-Abl activity from c-Src activity, but not compatible from incompatible peptides. (G) In canonical PTKs, the Asp of the HRD motif initiates catalysis by deprotonating the tyrosine hydroxyl to enable nucleophilic attack of the γ-phosphate ^49,50^. (H) D_1_, the distance between γ-phosphate and the oxygen on the central tyrosine (Y_0_, standard conformation; red highlight in G). (I) F1, the percent of the trajectories where the tyrosine (Y_0_; blue highlight in G) hydrogen bonds with the Asp122/Asp120 (c-Abl/c-Src). Error bars reflect the standard error of the mean (n = 3).

### Computational tools struggle to predict substrate orthogonality

The sensitivity of c-Src to the -1His mutation in the c-Src consensus motif is surprising, given its neutral enrichment in the PSSM (Fig. 1C). After all, +1Met was strongly depleted but had no effect on c-Src compatibility with the c-Abl consensus (Figs. 1C and 2C). We used molecular modeling to dive deeper. We began by using AlphaFold 3.0 (AF3)^45^ to predict PTK-substrate complexes; although AF3 is biased by its training data^46^, which contains only compatible PTK-peptide interactions, it provides a reasonable starting point for molecular dynamics (MD) simulations^47,48^. As expected, both PTKs exhibited similar catalytically competent conformations for both peptides (Figs. 5B and S22). We followed up by using MD to examine PTK-substrate complex stability. Interestingly, although the RMSDs of peptide backbones showed an initial shift for several peptides—a slight readjustment in bound pose—and this shift was largest for the -1His peptide bound to c-Src, they were small and stable for all four complexes, suggesting stable binding (Fig. 5D-5E). When we combined snapshots from all four sets of MD trajectories and sorted the peptide backbone coordinates into two clusters, the clusters grouped mainly by enzyme rather than substrate compatibility (Figs. 5F, S24).

In canonical PTKs, the Asp of the HRD motif initiates phosphorylation by deprotonating the tyrosine hydroxyl of the substrate to support nucleophilic attack of the nearby γ-phosphate (Fig. 5G; ^49,50^). To check for differences in catalytic competency between complexes, we measured (i) D_1_, the average distance between the γ-phosphate and the oxygen on the central Tyr (Fig. 5H), and (ii) F_1_, the frequency of contacts between the catalytic Asp and the same Tyr (Fig. 5I); here, we examined H-bonding between the two sidechains^51^. The bound poses did not show significant differences in either D_1_ or F_1_. Ultimately, the MD simulations do not indicate that the -1His peptide binds unstably to c-Src and suggest that the outsized influence of the -1His mutation may result from subtle effects that require advanced methods (e.g., longer simulations or enhanced sampling) to resolve clearly.

### Dual screening reveals peptides with surprisingly orthogonal PTK compatibilities

The inability of molecular modeling tools and PSSM maps to predict the impact of the - 1His mutation on c-Src compatibility is striking and illustrates the potential for minor sequence differences to cause large, functionally influential shifts in PTK-substrate interactions. To search for substrates that might shift specificity toward c-Src over c-Abl, we added a second screening step that selects for PTK-incompatible peptides. This step uses a “decoy substrate” consisting of a peptide fused to green fluorescent protein (sfGFP). When phosphorylated, the decoy binds to the SH2 domain without recruiting RNA polymerase; when screened in the presence of a functional B2H system (i.e., a peptide capable of localizing RNA polymerase to the GOI), only PTK-incompatible decoys (i.e., those that fail to interfere) permit survival (Fig. 6A).

**Figure 6.**
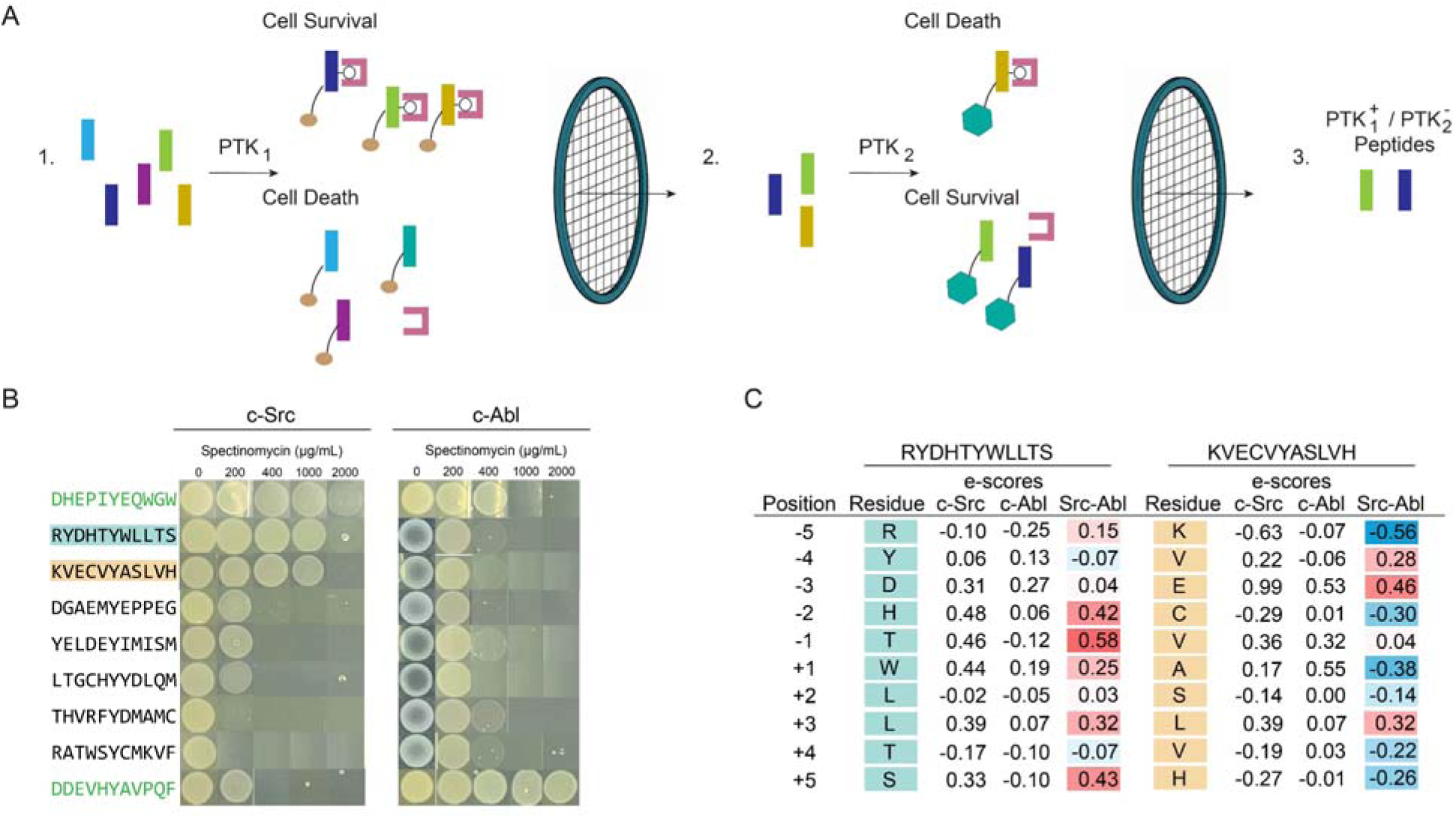
A dual-selection screen identifies substrates with unexpectedly orthogonal PTK compatibilities. (A) Screen for substrates that are compatible with PTK_1_ but not PTK_2_. In step 1, we use our standard B2H to screen a degenerate peptide library for substrates that are compatible with PTK_1_ (c-Src). In step 2, we attach the substrates selected in step 1 to a decoy (i.e., GFP) and screen the resulting decoy library in the presence of a functional B2H system for PTK_2_ (c-Abl). Only incompatible substrates, which do not sequester SH2 domains, permit B2H function and antibiotic resistance. (B) B2H-based assessment of a handful of substrates discovered in our two-step screen. We identified two hits (cyan and orange). References: c-Src consensus -1Thr (top, green) and c-Abl consensus -1His (bottom, green) peptides from Fig. 3D. (C) PSSM-based enrichment scores of each amino acid in two orthogonal substrates highlighted in B. Data depict (B) images of n = 3 biological replicates (Fig. S19) of (C) averages of n = 2 biological replicates.

To search for substrates compatible with c-Src but not c-Abl, we took peptides screened for c-Src compatibility using our original method, fused them to the decoy, and screened the decoy library in the presence of a functional B2H system for c-Abl (i.e., a B2H with c-Abl and its consensus peptide fused to rpoZ). After picking a handful of colonies, we identified two highly orthogonal substrates (Figs. 6B and S25). Their sequences were surprising. In general, their constituent residues had mild enrichment or depletion values (i.e, -1 to 1), relative to those observed in consensus motifs, and enrichment differences between the PTKs were minor (Fig. 6C). For one substrate, most amino acids were slightly more enriched for c-Src than for c-Abl; for the other, they were slightly more depleted. The PTK-selective interactions exhibited by these substrates are difficult to infer from PSSMs or previously developed substrate compatibility scores (Table S2). They provide a powerful demonstration of context-dependent amino acid preferences and highlight the extraordinary nuance of intrinsic substrate specificities.

## DISCUSSION

Protein tyrosine kinases (PTKs) regulate cellular biochemistry by phosphorylating distinct subsets of proteins. Precise knowledge of their substrate specificities is important for understanding signaling in healthy cells and how it is disrupted in disease. In this study, we developed a facile screening system for substrate specificity and used it to explore the limitations imposed by a common—sometimes implicit—assumption that amino acid preferences at each position within a peptide are independent of its sequence. This assumption has been acknowledged as imperfect in prior demonstrations of sequence-dependent preferences^22,29^, but the extent of that imperfection remains poorly understood.

Our B2H systems generated PSSMs consistent with previous screening methods and enabled rapid comparisons of specific PTK-substrate pairs via bacterial growth experiments. Our PSSMs gave functional consensus motifs, each of which was most compatible with its cognate kinase, but obscured differences in promiscuity between PTKs. The -1 position and C-terminal region were particularly important for controlling PTK compatibility. In follow-up analyses, we identified mutations within these sites that could improve inter-PTK orthogonality or confer sensitivity to a substrate length. Each effect was not readily predictable from our base PSSMs, where the implications of specific enrichment values for substrate compatibility appeared to differ by site, substrate sequence, and PTK.

Recent advances in protein modeling suggest that enzyme-substrate compatibility might one day be predictable from sequence data alone^52–54^. Unfortunately, peptides sample a large conformational space that is poorly represented in the Protein Data Bank,^55,56^ a flexibility that complicates the use of both AI/ML-based methods and MD simulations to predict binding poses. The dynamics of PTKs, which rearrange during binding and catalysis^57,58^, complicate matters further. Our studies of a substrate and a mutant are illustrative. Using our B2H-based assay, we identified a substrate for which a single mutation shifted its compatibility from two PTKs to one. Intriguingly, both AF3 and replicate simulations were unable to find prominent differences in PTK-substrate interactions. As with PSSMs, molecular modeling struggled to predict the impact of a single amino acid substitution on compatibility.

High-throughput methods to profile the average substrate specificity of PTKs are myriad, but complementary methods to characterize other context-dependent features of these interactions are rare. We developed a dual selection system that uses sequential screens of functional and decoy substrates to identify peptides that are compatible with one PTK but not another. This method produced substrates that were compatible with c-Src, but not c-Abl—an orthogonality that was not obvious from their sequences or our PSSMs. These substrates highlight the pronounced nonlinearity of PTK-substrate interactions. In future work, similar methods could yield “orthogonality maps” that show differences in the positional sensitivities of closely related PTKs. Detailed knowledge of the molecular basis of orthogonality in PTK-substrate interactions could help resolve the contributions of functionally related PTKs to complex signaling cascades or guide the design of PTK-specific sensors, probes, or therapeutics.

## MATERIALS AND METHODS

Details of cell strains, cloning and molecular biology, substrate screening, drop-based assays, NGS sequencing and analysis, quantitative comparisons of substrates, kinetic assays, protein and initial structure analysis, molecular dynamics simulations, trajectory processing and analysis, statistical analysis are provided in *SI Appendix, Materials and Methods*.

### Data, Materials, and Software Availability

Materials, supplementary figures and supplementary tables are provided in *SI Appendix, Materials and Methods.* Enrichment scores, cosine similarity scores, and kinetic data are provided in supplementary tables S8, S9, and S10. GROMACS input files used for trajectory processing and analysis are available at https://github.com/annette-thompson/PTKs_2026

## SUPPLEMENTARY MATERIAL

Supplementary figures detailing position-specific scoring matrices (PSSMs) and deep mutational scanning (DMS) data; comparisons of biological replicates, PSSM and DMS data, and PSSMs from this and other studies; growth-based assays of antibiotic resistance afforded by different B2H systems; visual representations of cosine similarity calculations and related heatmaps; plots of position-specific enrichment averages for each PTK; a standard curve for in vitro kinetic measurements of PTK activity on peptide substrates; and Tables of gene sources, plasmids, primers, library statistics, and data from all figures.

## ACKNOWLEDGEMENTS

This work was supported by funds provided by the National Institute of General Medical Sciences of the National Institutes of Health (H.E.A., A.T., and J.M.F., R35GM143089; M.R.S., R35GM158359) and the National Science Foundation (A.T., 2025391967). This work utilized the Alpine high-performance computing resource at the University of Colorado Boulder. Alpine is jointly funded by the University of Colorado Boulder, the University of Colorado Anschutz, and Colorado State University and with support from NSF grants OAC-2201538 and OAC-2322260.

## AUTHOR CONTRIBUTIONS

J.M.F. and H.E.A. conceived of research. H.E.A., N.O., J.M.K., and J.M.F. designed experiments. H.E.A. and N.O. carried out initial cloning of B2H systems. H.E. carried cloned select PTK-substrate pairs in follow-up analyses, and carried up selection assays, PSSM analyses, drop-based plating, and kinetic measurements. A.T. carried out molecular simulations with guidance from M.S. All authors analyzed data. H.E.A., A.T., and J.M.F. wrote the paper with comments from all authors.

