## Supporting Information for "Context-dependent peptide recognition shapes tyrosine kinase substrate specificity beyond consensus motifs"

**This PDF file includes:**

Supporting Information Materials and Methods

SI Notes

Figures S1-S25

Tables S1-S10

SI References

**Supporting Information Materials and Methods**

**Materials.** We purchased yeast extract, sodium chloride, potassium phosphate monobasic, potassium phosphate dibasic, tetracycline hydrochloride, agarose, and black non-binding flat-bottom plates from Fisher; spectinomycin dihydrochloride pentahydrate from MilliporeSigma; Phusion, DpnI, NdeI, XhoI, Taq DNA ligase, CutSmart buffer, Phusion High-Fidelity buffer, dNTPs, T4 Polynucleotide Kinase, Nt.BbvCI, Nb.BbvCI, Exonuclease III, Exonuclease I, NAD+, DTT, ATP, and T4 DNA ligase from New England Biolabs; agar from Becton Dickinson; tryptone from Research Products International; glycerol from Macron; kanamycin sulfate from IBI Scientific; carbenicillin from Gemini Bioproducts; ATP Gold and ADP Quest kit from Eurofins DiscoverX Products; primers from Integrated DNA Technology, kinases from CarnaBio; Zymo Clean & Concentrator kit and E.Z.N.A Plasmid Mini Kit I from VWR; and custom peptides (DDEVIYAVPQF, DDEVHYAVPQF) from the CU Boulder Peptide Synthesis Facility.

***E. coli* Strains.** We used chemically competent NEB Turbo (#C2984H) cells for molecular cloning and Fisher One Shot Top10 electrocompetent cells (#C404003) for library preparation, and electrocompetent and chemically competent S1030 cells (Addgene #105063) for drop plating. We generated chemically competent and electrocompetent cells with standard protocols (DMSO or glycerol and washing).[1]

**Cloning and Molecular Biology.** We used Gibson assembly (50°C for 1 hour) to construct all plasmid templates. Table S3 lists gene sources; Table S4 describes each plasmid; and Table S5 lists primers for plasmid assembly and NGS sequencing. To prepare substrate libraries for B2H screening, we use used degenerate primers to randomize the substrate sequence in a DNA amplification reaction (98°C 2 min; 25x of 98°C 20 seconds, 65°C 15 seconds Δ0.5°C, 72°C 30 seconds/kbp; 10x 98°C 20 seconds, 55°C 15 seconds, 72°C 30 seconds/kbp; 72°C 2 minutes), NdeI and XhoI restriction enzymes to prepare the amplification product (i.e., $X_5$-Y-$X_5$ with DNA overhangs) for re-insertion into a pMM532 plasmid harboring substrate-$Rp_\omega$, and a T4 DNA ligase to ligate the substrate and plasmid. Our deep mutational scanning approach followed the protocol described by Wrenbeck et al.[2] We confirmed substrate diversity by checking a small number of plasmids with Sanger Sequencing (>50% of plasmids carried a unique, non-template, or $S_{Y/F}$, substrate) or by sending the library for whole-plasmid sequencing to ensure >50% sequences differ from the template peptide (Quintara).

**Substrate Screening.** We screened substrates for compatibility with PTK-specific B2H systems by measuring their enrichment on plates containing high concentrations of spectinomycin. In brief, we dialyzed plasmids containing our substrate libraries for 2 hours (MF-Millipore 0.025 µm MCE membrane filters, 25mm), electroporated ~1000 ng of DNA into s1030 cells encoded with a PTK-specific B2H system, recovered cells for 1 hour (37°C shaker), and plated them onto LB agar plates (5 g/L yeast extract, 10 g/L tryptone, 10 g/L sodium chloride, 20 g/L agar, pH=7.5, 50 µg/mL kanamycin, 50 µg/mL carbenicillin, and where necessary, 10µg/mL tetracycline) with and without spectinomycin (0 µg/mL and a selection condition depending on the PTK). For each PTK, we chose our "selection" condition (i.e., spectinomycin concentration) by plating with substrates S (EPQ**Y**EEIPIYL) and $S_{Y/F}$ (EPQ**F**EEIPIYL) on varying concentrations of spectinomycin and selecting the highest concentration at which (i) S gave many colonies and (ii) $S_{Y/F}$ gave none. These sequences correspond to positive and negative controls from our previous work with the B2H system.[3,4] In some cases, we repeated screens at lower spectinomycin concentrations to improve reproducibility between replicates.

We adjusted screening conditions for each kinase. For c-Src, c-Abl, EPHA2, and FGFR2, we incubated reference and selection plates (0 and either 600, 200, 200, or 100 µg/mL spectinomycin, respectively) for 16-18 hours at 37°C. For JAK2, we incubated reference and selection plates for 48 hours at 30°C, which yielded a higher signal than 37°C. For the decoy library, our control sequences (DDEVHYAVPQF and DDEEDYEEIWW) suggested 100 µg/mL as an ideal screening condition. For all screens, we pooled cells from select plates, extracted plasmid DNA (E.Z.N.A. Plasmid Mini Kit I, Omega Bio-Tek), amplified substrate pools with primers listed in Table S5 (98°C 2 min; 15x of 98°C 20 seconds, 55°C 15 seconds, 72°C 12 seconds; 72°C 2 minutes), gel purified the amplicons, and sequenced amplicons with next-generation sequencing (AmpExpress, Quintara). Estimated library sizes in Table S6.

**Drop-based Assay.** We measured the antibiotic resistance afforded by B2H systems that differ by a single component to assess differences in PTK-substrate compatibility. To begin, we used heat shock to transform s1030 cells with plasmid-borne B2H components, recovered cells for 1 hour (37°C shaker), plated cells on LB agar (as above), and incubated them for 16-18 hours (37°C). We used 3 colonies from each plate to inoculate 1 mL of liquid TB (12 g/L tryptone, 24 g/L yeast extract, 2.28 g/L potassium phosphate monobasic, 12.5 g/L potassium phosphate dibasic, 2% v/v glycerol) in 96 well-blocks, which we incubated in an incubator shaker (37°C at 225 rpm) overnight. We diluted overnight cultures to an $OD_{600}$ = 0.1 in liquid TB (as above), dropped 3.5 µL onto LB agar plates with varying concentrations of spectinomycin (0, 200, 400, 600, 800, 1000, 1200, 2000, and 3000 µg/mL), and incubated plates for 16-18 hours at 37°C prior to photographing. For Jak2, we altered this procedure by incubating plates for 24 hours at 30°C. For all PTKs, we used S and $S_{Y/F}$ as controls to get a rough sense of the resistance afforded by compatible (S) and incompatible ($S_{Y/F}$) substrates.

**Next-Generation Sequencing (NGS) and analysis.** We processed and analyzed NGS data using BBMap, a short-read aligner, and our own code. We cleaned Fastq files from Quintara using the BBMap short read aligner [5], merged forward and reverse read files for each sample, and selected only reads with a quality score of 20 (99% accuracy for bases). Final reads include (i) sequences from the reference plates and (ii) sequences from the selection plates. We processed these groups with a Python script that extracts the substrate sequences and counts the number of residues at each position. We used these counts to calculate enrichment by using Eq. 1, where $A_{ij}$ is the read

$$E = log_2\left(\frac{A_{ij}}{A_{tot}}\right) - log_2\left(\frac{R_{ij}}{R_{tot}}\right) \qquad (Eq. 1)$$

count of amino acid $i$ (e.g., Ala) at position $j$ (e.g., -1) in the substrate sequences from the selection sample, $A_{tot}$ is the total number of reads in the selection sample, $R_{ij}$ is the read count of amino acid $i$ at position $j$ from the reference sample, and $R_{tot}$ is the total number of reads in the reference sample. Table S7 contains consensus sequences derived from average enrichment scores; Table S8 provides the individual and average enrichment scores for two biological replicates for each of our screens; our heatmaps show average enrichment scores. The inclusion of stop codons in our determination of consensus sequences did not affect PTK compatibility, but "secondary consensus" peptides constructed with the 2$^{nd}$ most enriched residue at each position in a PSSM resulted in poorly compatible peptides (Fig. S20). We also checked our c-Src consensus peptide, a previous c-Src consensus peptide,[6] the S (MidT peptide) sequence, and a previously reported c-SH2 consensus peptide[6] against our c-Src PSSMs and the previously reported c-Src PSSM[6] (Fig. S21). We found that both consensus peptides contained residues enriched for both datasets, and the poorly compatible S peptide less so. We included the c-SH2 peptide to check for bias but did not see much influence from it.

**Quantitative comparisons of substrate preferences.** We used two metrics to compare the substrate preferences of PTKs. First, we calculated the cosine similarity ($S_{ij}$) of enrichment patterns for each binary combination of PTKs (Fig. S6). In Eq. 1, $M_i$ and $M_j$ are vectorized

$$S_{ij} = \frac{M_i \cdot M_j}{\|M_i\| \|M_j\|} \qquad \text{(Eq. 2)}$$

enrichment maps for PTKs $i$ and $j$, and $S_{ij}$ is a metric for the similarity of those maps. As described in the main text, this metric weights all positions equally and assumes symmetric preferences between PTKs. Next, we calculated the cosine similarity ($P_{ij}$) of enrichment values for a specific substrate sequence. In Eq. 2, $l_i$ is a vector of the enrichment values each amino acid

$$P_{ij} = \frac{l_i \cdot l_j}{\|l_i\| \|l_j\|} \qquad \text{(Eq. 3)}$$

in the consensus motif for $PTK_i$; $l_j$ is a vector of enrichment values for the same sequence for $PTK_j$; and $P_{ij}$ describes the compatibility of the consensus motif for $PTK_i$ with $PTK_j$. Table S9 contains all cosine similarity scores.

**Kinetic assays.** We measured PTK activity on peptides by using a continuous fluorescence-based assay (ADP Quest, Eurofins). In this assay, PTK-mediated phosphorylation of peptides converts ATP to ADP; pyruvate kinase converts ADP and phosphoenolpyruvate to ATP and pyruvate; pyruvate oxidase decarboxylates this newly generated pyruvate to produce hydrogen peroxide ($H_2O_2$); and peroxidase uses this $H_2O_2$ to oxidize Amplex Red to resorufin, a highly fluorescent compound. We used the buffer supplied with the kit to prepare solutions of purified peptides (>98%) and PTKs, which we used to prepare reactions with 10 nM c-Abl or 20 nM c-Src, 100 μM ATP, and 0-400 μM peptide. We monitored reaction progress with a plate reader ($\lambda_{ex}$ = 530, $\lambda_{em}$ = 590 nm, every 9 seconds, room temperature), used a standard curve to convert fluorescence to ADP concentration (Fig. S16), and subtracted initial rates of samples without kinase to account for background ATP hydrolysis. We plotted rates as a function of substrate concentration to estimate Michaelis-Menten parameters ($k_{cat}$ and $K_M$) using DataGraph. Table S10 contains data from our in vitro kinetic assays.

**Protein structure analysis.** We used AlphaFold 3.0[7,8] (AF3) to predict four PTK-substrate complexes: (i) c-Abl and DDEVIYAVPQF, the c-Abl consensus peptide, (ii) c-Abl and DDEVHYAVPQF, the -1His mutant, (iii) c-Src and the consensus, and (iv) c-Src and the mutant. For these analyses, we used PTK sequences of c-Src$_{270\text{-}523}$ and c-Abl$_{242\text{-}493}$. Each complex included ATP and $Mg^{2+}$; for c-Src, we included pTyr at 419 on the activation loop (a post-translational modification that is important for activation and substrate recognition).[9] We compared these structures to PDB structures 1Y57,[10] 1M52,[11] and 2G2I[12] to check for active conformation in predicted structures (Fig. S22).

**Initial structural analysis.** To initialize MD simulations, we selected eight PTK-substrate complexes comprised of two groups: (i) AF3 predictions of c-Abl and c-Src bound to the consensus peptide or its -1His mutant and (ii) crystal structures of c-Abl (PDB 3KF4) and c-Src (PDB 1YI6) overlaid with the same AF3-derived peptide conformations. After surveying the PDB, we selected crystal structures that showed an active DFG conformation and did not clash with ATP or the bound peptide, and we filled in missing residues with MODELLER. We selected AF3 structures, in turn, by selecting complexes with the lowest RMSDs relative to the crystal structures (Fig. S22).

For all PTK-peptide pairs, we used three independent sets of energy minimization and equilibration steps to initialize replicate simulations, which were then run for 500 ns. We visually inspected simulations using VMD 1.9.4 to confirm peptide retention. In general, trajectories for the AF3 structures exhibited lower protein backbone RMSDs to their centroid structures than those seeded from crystal structures with predicted peptide poses (Fig. S23), so we used AF3 trajectories for subsequent analyses. We note: In our initial comparison of AF3 and crystal structures, we used a Y419E mutant of Src—a mimic of the activating pTyr that is easy to model [13,14]; after confirming the enhanced stability of the AF3 structures for c-Src and c-Abl, we used the pTyr variant of c-Src for our subsequent analyses (Fig. 5 D-I).

**Molecular dynamics simulations.** We carried out MD simulations using structures from the PDB and AF3. We assigned protonation states to all species at pH 7.5 using pdb2pqr, generated force field parameters using the OpenFF Interchange, and parameterized proteins and peptides using the AMBER14SB force field. We assigned ATP partial charges using NAGL AM1-BCC GNN model (v0.1.0-rc3), followed by parameter assignment with the OpenFF Sage 2.2.1 force field, and added parameters for magnesium ions to the OpenFF framework by using values from the AMBER force field. For all complexes, we used a TIP3P water box (at least 10 Å to the periodic boundary) and neutralized charges with $Na^+$ ions (final salt concentration of 0.15 M).

We carried out MD simulations with GROMACS 2022.4 on the Alpine high-performance computing cluster at the University of Colorado Boulder. For each of three replicates, we carried out energy minimization until the maximum force fell below 500 kJ/mol/nm, followed by NVT equilibration at 300 K for 100 ps and NPT equilibration at 300 K and 1 atm for 100 ps. For all simulations, we used a velocity-rescaling thermostat[15] and a stochastic cell-rescaling barostat[16], and we applied hydrogen mass repartitioning (H mass of 3 amu) to enable long integration timesteps (3 fs in the final production run). For each PTK-substrate pair, we carried out three replicate simulations for 500 ns.

**Trajectory processing and analysis.** We removed periodic boundary condition artifacts (GROMACS tools) and truncated trajectories to equilibrated regions (backbone RMSD fluctuations of less than 0.1 nm over 10% of the trajectory and mean RMSD changes of less than 0.05 nm over 1% of the trajectory, evaluated using 100 ps windows and rounding the final equilibration window to the nearest 5 ns). To analyze protein backbones, we truncated proteins to conserved regions shared by c-Abl and c-Src (i.e., the KLG and QAF motifs, and excluding disordered terminal regions) and calculated RMSDs relative to the centroid of the largest structural cluster found using gmx rms. As we mention above, the RMSDs of runs seeded with AF3 structures were more stable than those seeded with PDB crystal structures (Fig. S23), so we focused subsequent analyses on the AF3-initialized trajectories. To analyze peptide backbones, we calculated RMSDs relative to the initial starting structure after aligning each trajectory frame to the protein backbone using MDAnalysis.

We calculated inter-atomic distances and contacts using MDAnalysis and identified hydrogen bonds using the geometric hydrogen-bond criteria implemented in MDAnalysis. We performed k-means clustering as implemented in scikit-learn, with a maximum of 1000 iterations. We obtained centroid structures of peptide backbones using k-means clustering with a single cluster (n_clusters = 1) and visualized them with PyMOL. All GROMACS input files

(.gro, .top, .mdp) are available in the associated GitHub repository (https://github.com/annette-thompson/PTKs_2026).

**Statistical analysis and reproducibility.** We measured kinetic data with 3-5 replicates per datapoint and calculated $k_{cat}$ and $K_M$ values with MATLAB by fitting data to a Michaelis-Menten curve (MATLAB, Table S10).

**SI Note 1. Peptide score.** In their work using bacterial peptide display (BPD) to profile PTK substrate specificity, Li et al.[6] proposed a method for using PSSMs to predict the PTK-peptide compatibility. In brief, they scored the compatibility of a specified peptide with a specific PTK by (i) summing the $\log_2$-fold enrichment of each peptide residue, (ii) dividing by the number of variable positions (e.g., 10 for most libraries), and (iii) normalizing such that the lowest and highest-scoring peptides were 0 and 1, respectively, for each PTK. This method, which gives the highest-scoring peptide (e.g., the consensus) a score of 1, assumes that amino acid preferences at each position in a peptide are independent of the sequences in which they occur.

### SI Figures

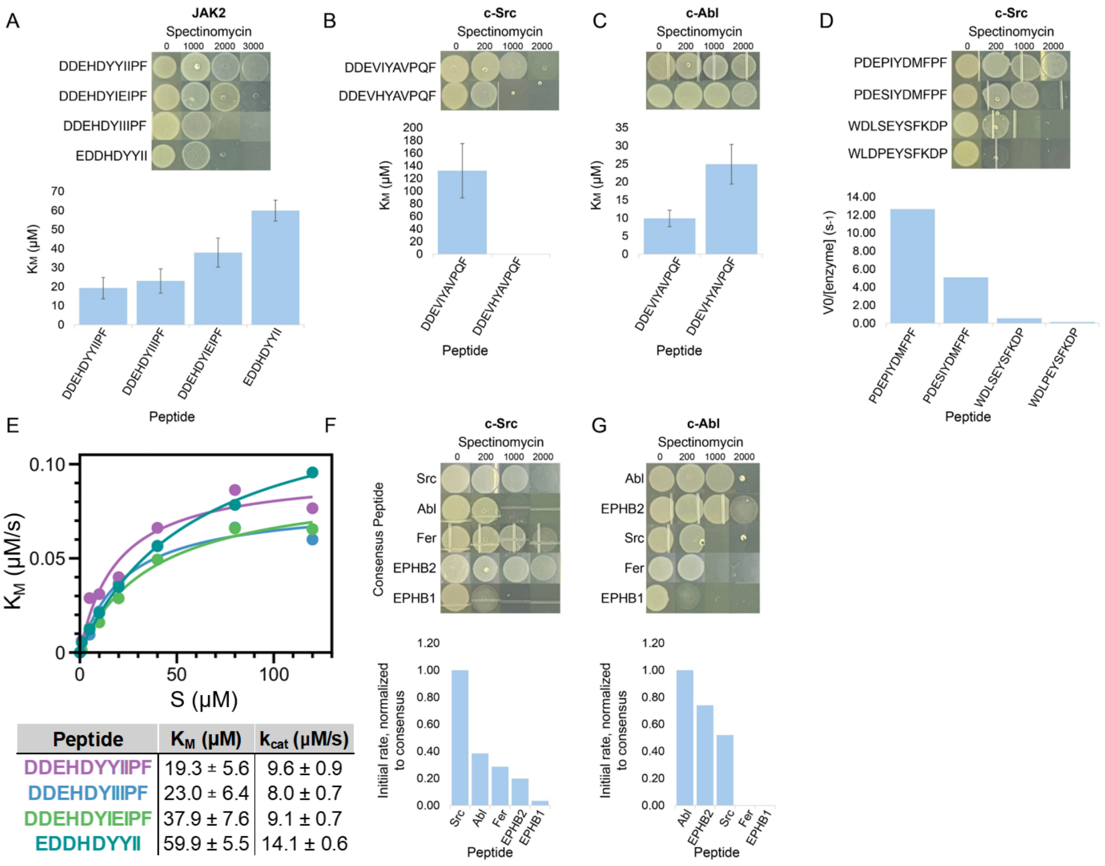


**Figure S1. PTK-peptide compatibility in drop assays and kinetic measurements across two papers.** (A) Drops and $K_M$ data for JAK2 kinase and four peptides. Drops for the +2Glu and +1Tyr both improve survival on spectinomycin as compared to the JAK2 consensus peptide, but There is no statistically significant difference between the three. However, the +1Tyr mutation does yield a different $K_M$ in the 11mer vs. 9mer consensus peptides. (B) Drops and $K_M$ for Abl consensus and Abl -1His mutant peptides with c-Src kinase (see Fig. 5A for Michaelis-Menten curves). (C) Drops and $K_M$ for Abl consensus and Abl -1His mutant peptides with c-Abl kinase (see Fig. 5A for Michaelis-Menten curves). (D) Drops of four mutant peptides from Li et al. 2023 tested in our B2H system for c-Src kinase compatibility, as well as kinetic data collected by Li et al. 2023 for these peptides and c-Src kinase[17]. As rate/enzyme decreases, so does survival on antibiotic. (E) Michaelis-Menten curves for JAK2 kinase and the four peptides in (A). (F) We tested consensus peptides from Li et al. 2023 for five different PTKs against c-Src using our B2H system and compared to their initial rate data. (G) The same consensus peptides from Li et al. 2023 tested against c-Abl using our B2H system, and their initial rate data. While c-Src is more promiscuous, the B2H screen for c-Abl compatibility follows initial rate data well. Data represent averages of n=3-5 ($K_M$, $k_{cat}$) or n=3 (cell drops).

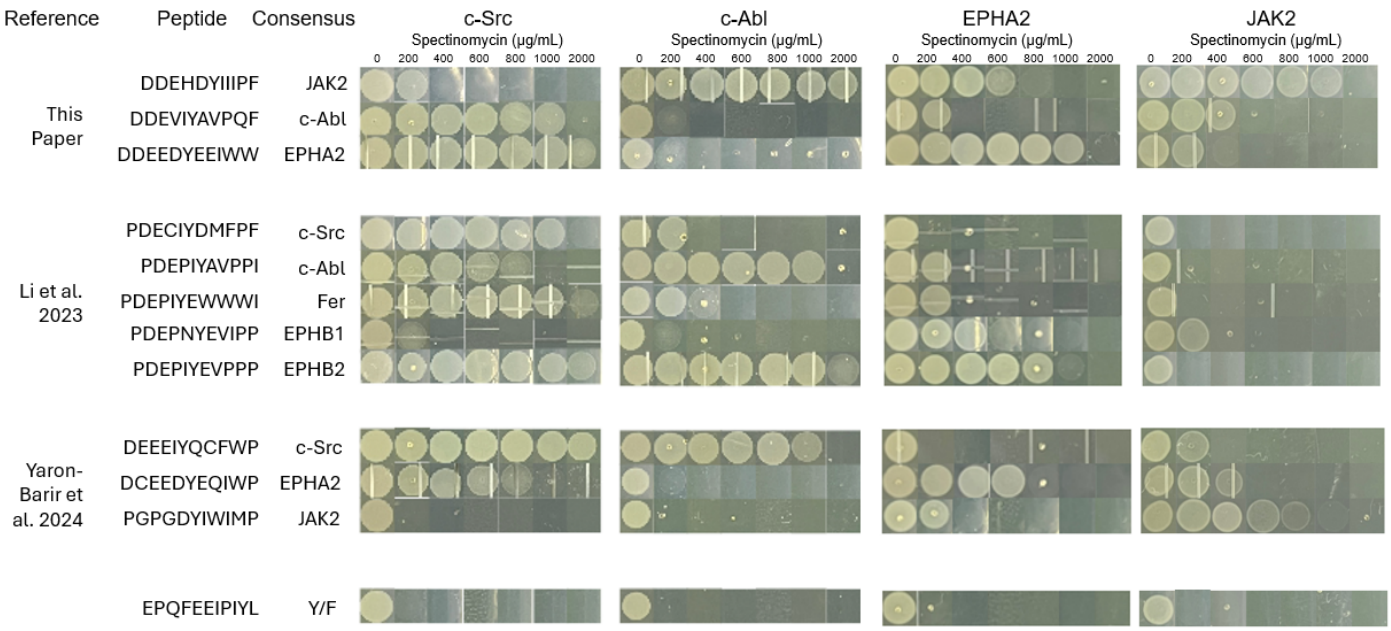


**Figure S2. Consensus peptides across three papers.** Consensus peptides from this paper, Li et al. 2023, and Yaron-Barir et al. 2024, tested for compatibility with c-Src, c-Abl, EPHA2, and JAK2 kinase using our B2H system. Plated at OD=0.1, 37C for 18 hours, n=3.

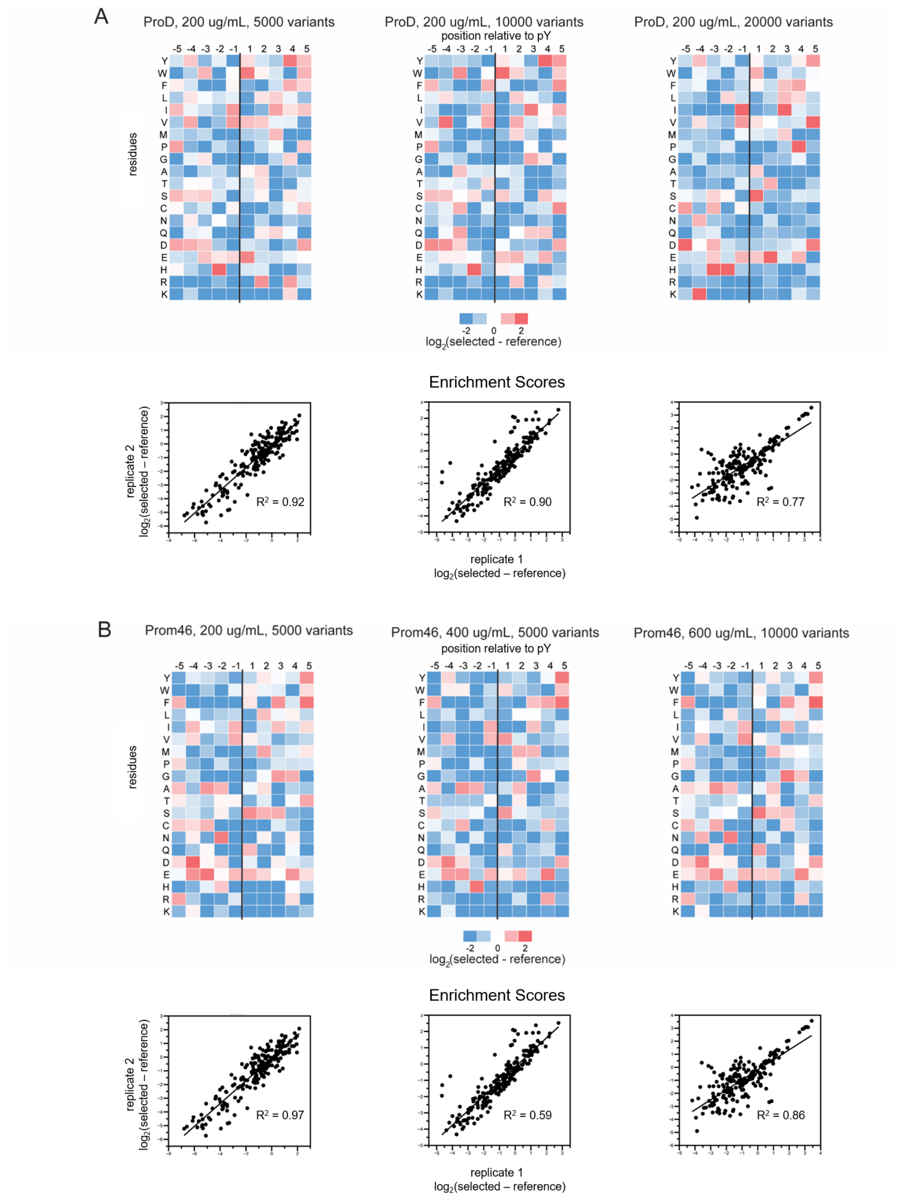


**Figure S3. Optimization of screening protocol.** (A) Increasing the size of the substrate library did not improve reproducibility between replicates (e.g., $R^2$ of enrichment). (B) Swapping a weak promoter (proD) for a strong promoter (prom46) enabled screening at high concentrations of spectinomycin (200, 400, and 600 µg/mL).

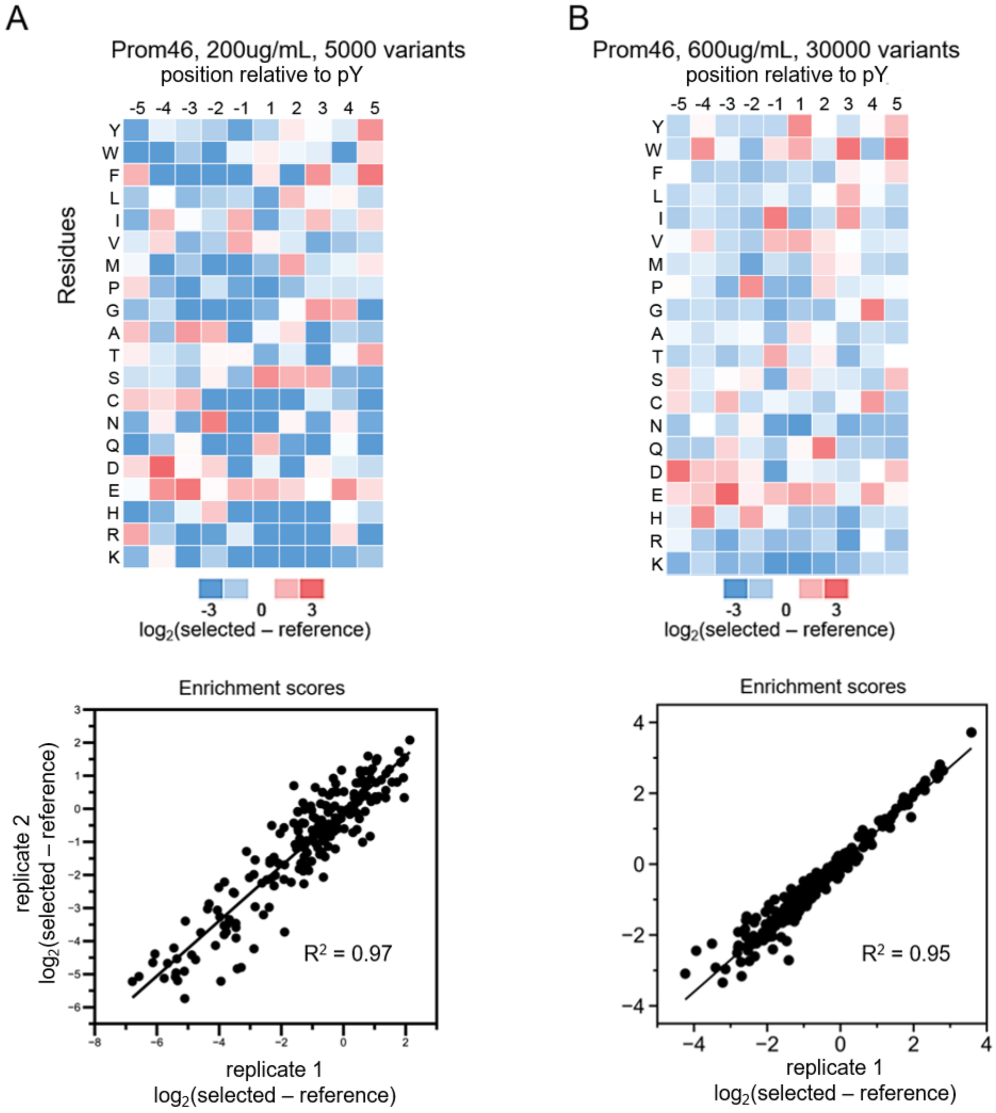


**Figure S4. Final screen parameters.** After optimization of our experimental protocol, our prom46-based system enabled good replicate agreement at (A) low and (B) high spec concentrations. We used the latter in our final screen (Fig. 1C).

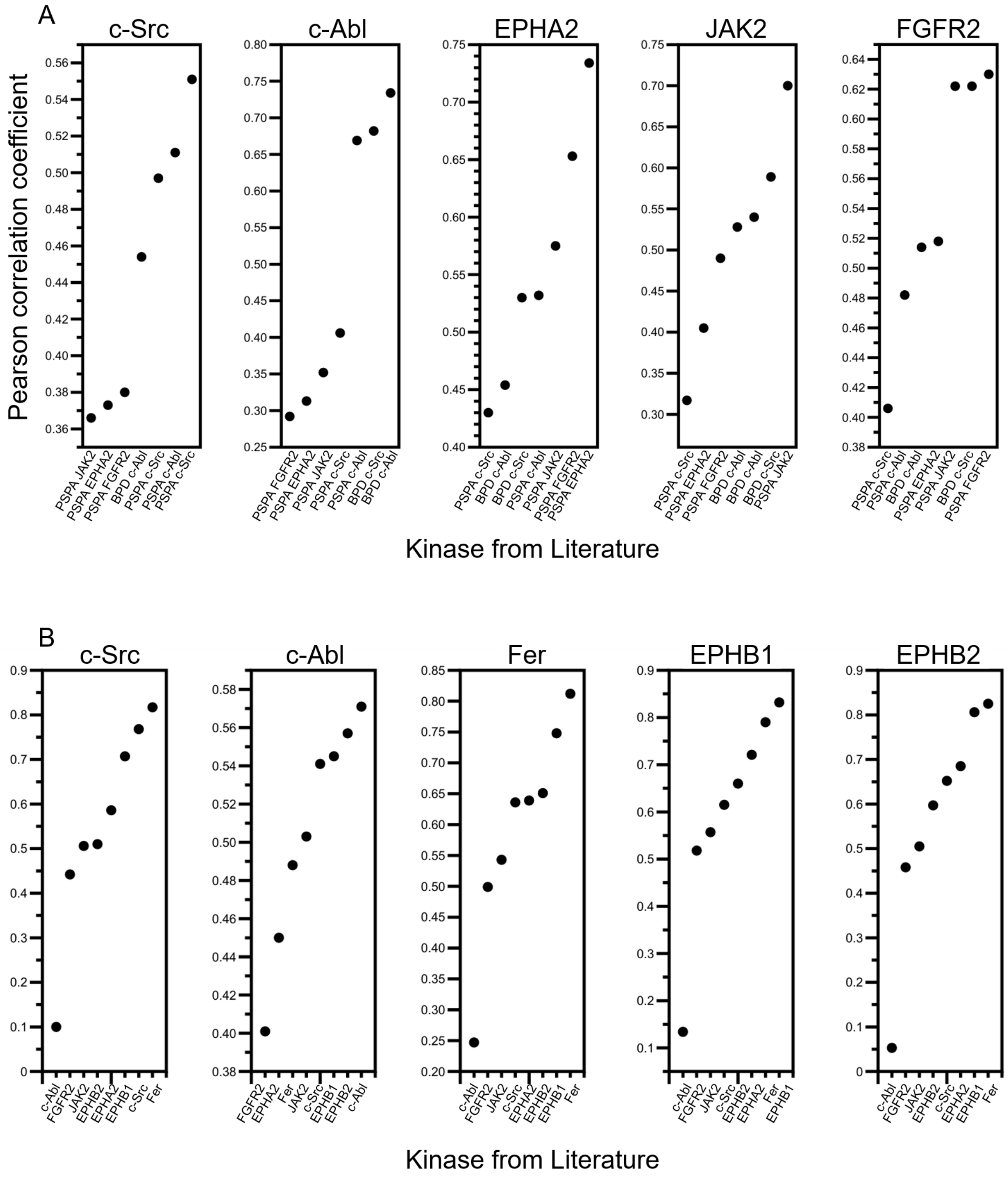


**Figure S5. Pearson correlation coefficients for PSSMs from this study and the literature.** We used Pearson correlation coefficients to compare methods. (A) Comparison of PSSMs generated by our method and those generated via (i) positional scanning peptide array (PSPA) [18]

or (ii) bacterial peptide display (BPD) [6]. (B) Comparison of PSSMs generated via BPD (each plot) and PSPA (X-axis). In general, the consensus preferences measured by our approach are as consistent with alternative methods (e.g., PSPA and BPD) as those methods are with each other. Each plot contains scores for a single BPD dataset.

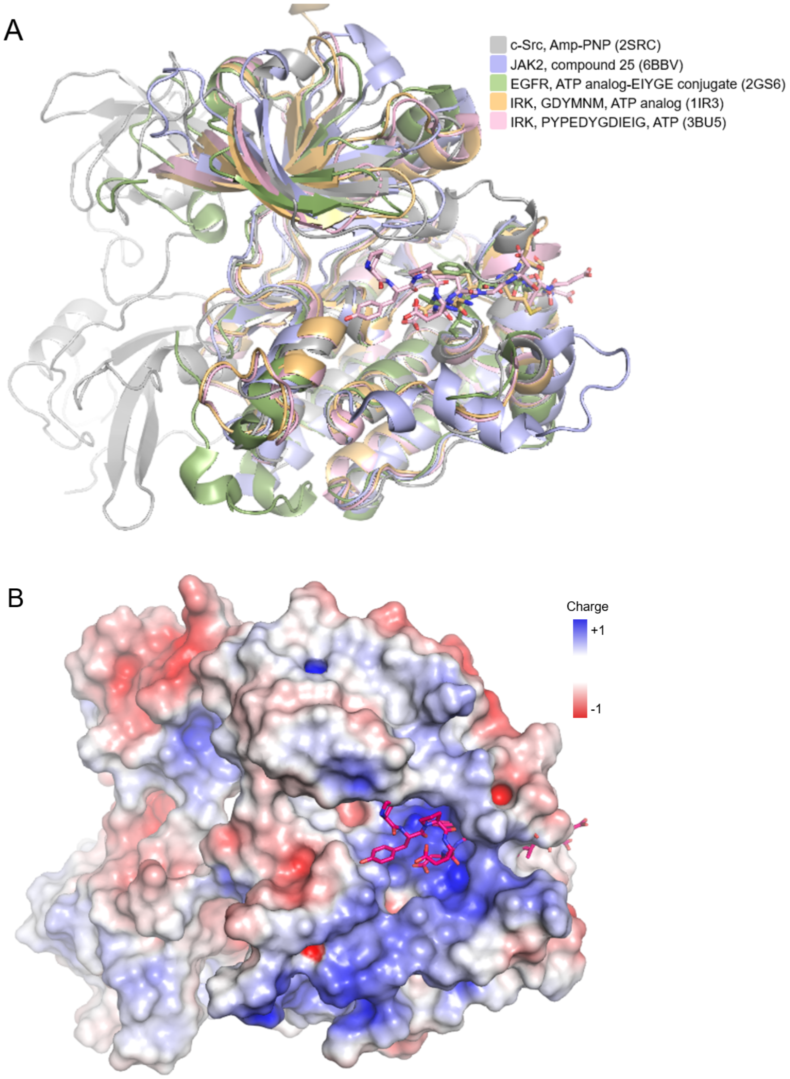


**Figure S6. A positively charged region in the substrate binding pocket.** (A) Alignment of c-Src kinase (PDB 2SRC, grey), JAK2 in complex with compound 25 (PDB 6BBV, blue), EGFR in complex with an ATP-peptide conjugate (PDB 2GS6, green), IRK in complex with IRS2 KRLB peptide (PDB 3BU5, pink), and IRK in complex with peptide substrate (PDB 1IR3, orange). The overlapping peptides and peptide analogues highlight the likely binding pocket for c-Src. (B) Electrostatic map of the c-Src kinase surface reveals a positively charged region in the putative peptide binding site (IRS2 KRLB peptide, from PDB 3BU5, in dark pink).

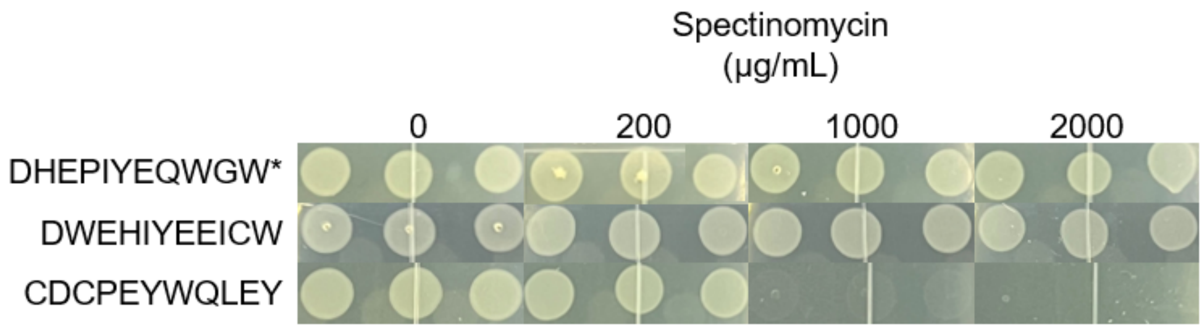


**Figure S7. Triplicate drop data of Fig. 1E.** Triplicate drops from Figure 1E. Drops plated at OD=0.1, grown 18 hours at 37ºC.

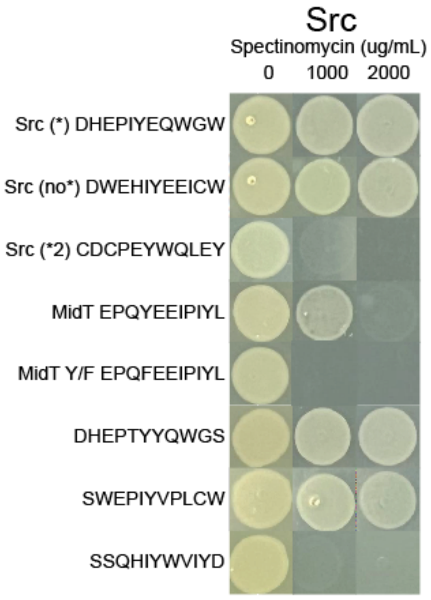


**Figure S8. Analysis of c-Src substrate preferences.** Consensus sequences for c-Src generated from heat maps with (*, DHEPIYEQWGW) and without stop codons (no *, DWEHIYEEICW) showed similar compatibilities in our growth-based assay. A substrate based on the 2nd most enriched residue at each position (*2, CDCPEYWQLEY) was not compatible. Highly enriched substrate sequences (DHEPTYYEWGS, SWEPIYVPLCW, SSQHIYVPLCW), which are reported in sequencing, show high compatibility in two out of three cases. Data from n=2.

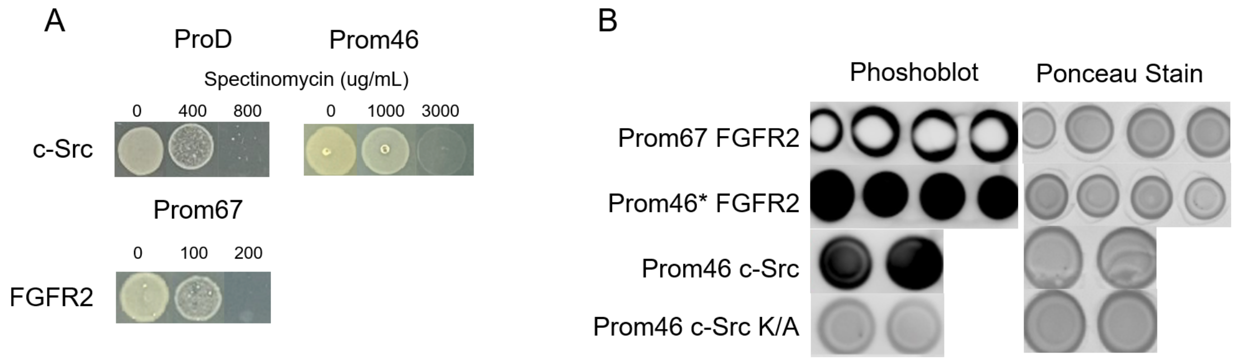


C

| Promoter | PTK | TIR | Phosphorylation | S-mediated spectinomycin survival (ug/mL) | Growth |
|---|---|---|---|---|---|
| prom46* | FGFR2 | 406.5 | high | N/A | slow |
| prom46 | FGFR2 | 241.8 | N/A | N/A | N/A |
| proD | FGFR2 | 482.6 | N/A | N/A | N/A |
| prom67 | FGFR2 | 196.4 | very high | 100 | normal |
| prom46 | c-Src | 215.4 | high | 3000 | normal |
| proD | c-Src | 430.0 | N/A | 800 | normal |

**Figure S9. Expression level variations in PTKs.** (A) Drop-based plating of *E. coli* harboring B2H systems for c-Src expressed with weak (proD) and strong (prom46) promoters. For c-Src, the strong promoter improves antibiotic resistance. For FGFR2, only a medium promoter (prom67) enabled growth. (B) Phosphoblots of *E. coli* harboring B2H systems for (top) FGFR2 expressed with strong (prom46*, a mutated prom46 that spontaneously arose during cloning) and medium (prom67) promoters, and (bottom) c-Src and c-Src K/A (prom46, controls). The phosphoblot suggests that prom46* may lead to toxic over-phosphorylation of the proteome. In A-B, Ponceau S serves as control by non-specifically staining proteins in reaction. (D) Summary of results from drop assays and phosphoblots. Prom46 FGFR2 could not be cloned, and proD FGFR2 was not tested. Placing a PTK under a promoter with a weaker predicted translation initiation rate (TIR) resulted in cells with normal growth rates and variable S-mediated spectinomycin survival, while preserving high phosphorylation. Too strong promoters may lead to toxicity by overphosphorylation by the PTKs being expressed. Predicted TIR rates calculated through the Salis Lab promoter calculator [19].

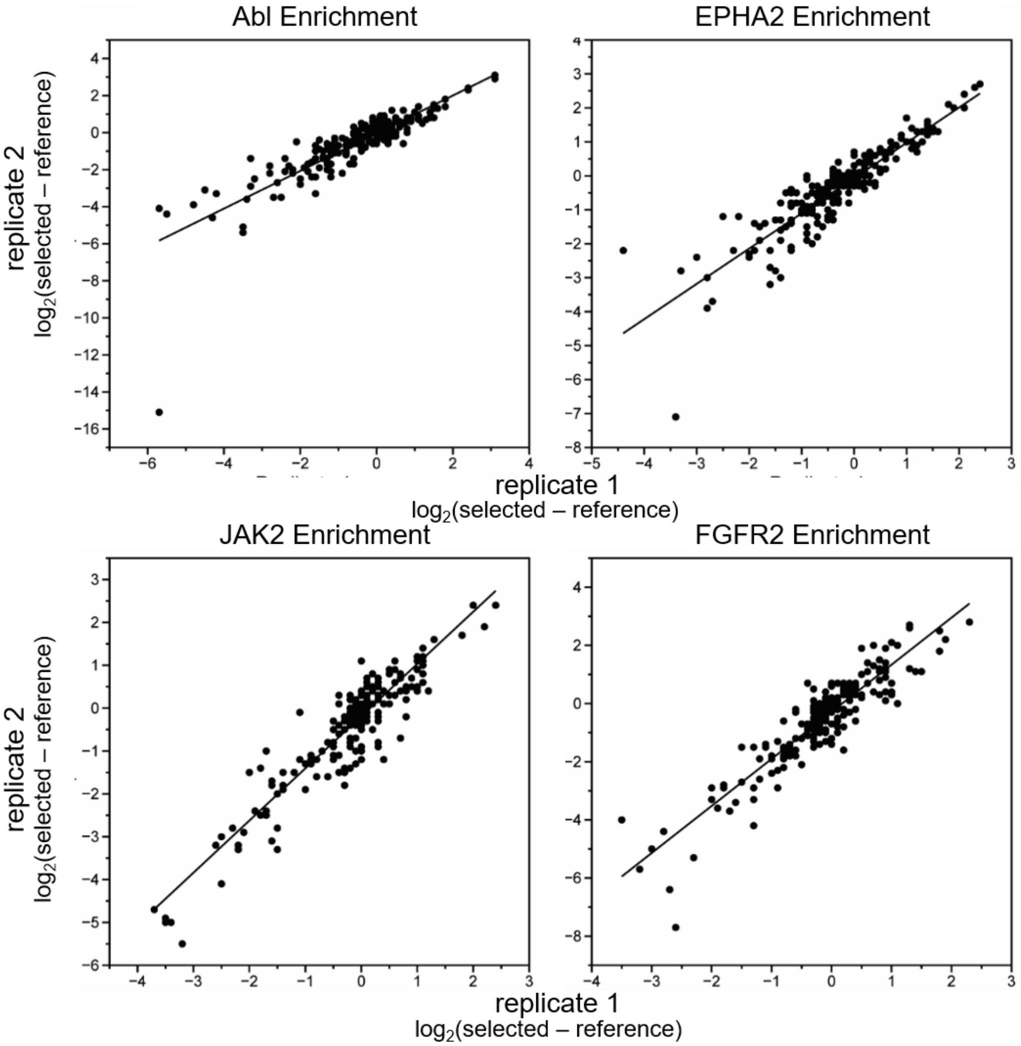


**Figure S10. Replicate data for heatmaps.** These data show the reproducibility (i.e., $R^2$ for biological replicates) of 11-residue PSSMs for the PTKs shown in Figure 2A. Summary: $R^2$ = 0.74 (c-Abl), 0.81 (EPHA2), 0.85 (JAK2), and 0.83 (FGFR2).

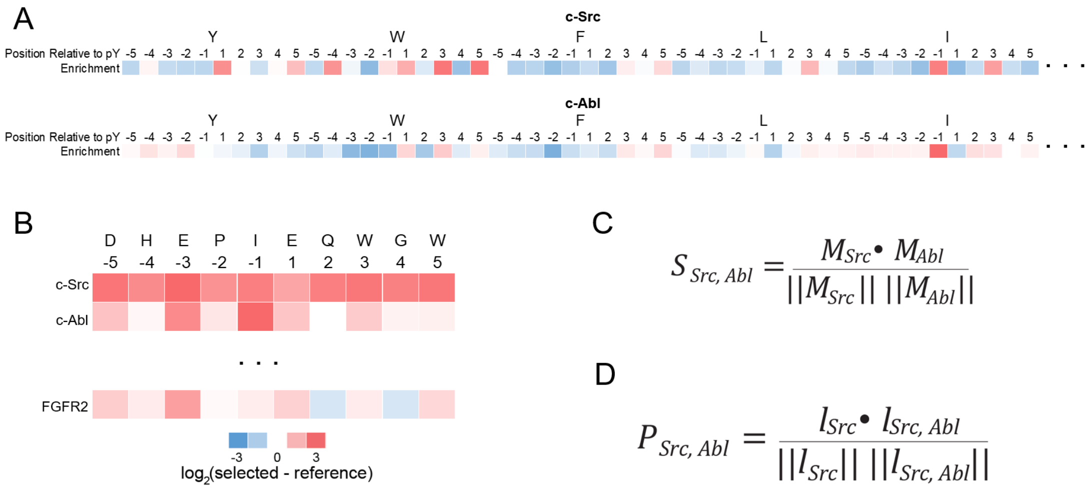


**Figure S11. Analysis of substrate preferences.** (A) To calculate the cosine similarity of two PSSMs (Fig. 2B), we vectorized their heatmaps. Pictured: c-Src and c-Abl PSSMs. (B) To compare the compatibility of a specific substrate with two PTKs (Fig. 2C), we used vectors of the substrate enrichment scores for each PTK. Pictured: the c-Src consensus sequence. (C-D) Exemplary calculations. (C) $S_{Src,Abl}$ is the cosine similarity of $M_{Src}$ and $M_{Abl}$, the PSSMs of c-Src and c-Abl. (D) $P_{Src,Abl}$ is the cosine similarity of $l_{Src}$, a vector of c-Src enrichment values for the c-Src consensus peptide, and $l_{Abl}$, a vector of c-Abl enrichment values for the same sequence.

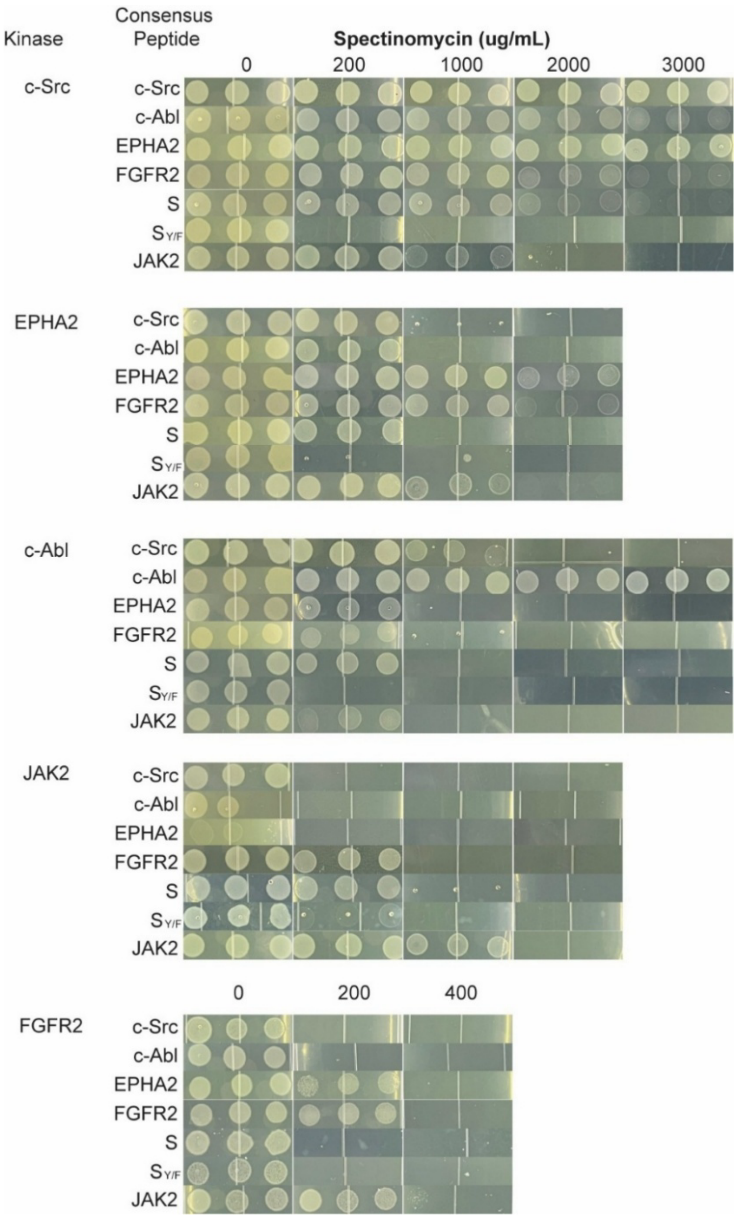


**Figure S12. Drop-based plating to examine PTK-substrate compatibility (Fig. 2).** Drop-based plating of s1030 cells harboring B2H systems for each of five PTKs ($B2H_{c\text{-}Src}$, $B2H_{c\text{-}Abl}$, $B2H_{EPHA2}$, $B2H_{JAK2}$, $B2H_{FGFR2}$), their consensus substrates, and substrate controls S and $S_{Y/F}$ ($pMM532_{c\text{-}Src}$, $pMM532_{c\text{-}Abl}$, $pMM532_{EPHA2}$, $pMM532_{JAK2}$, $pMM532_{S}$, and $pMM532_{S\_YF}$). Details: 3.5 uL drops of OD=0.1 cell culture placed on LB agar plates incubated at 37°C for 16-18 hours. For JAK2, we used 30°C and 24 hours. Images: drops for n = 3 biological replicates.

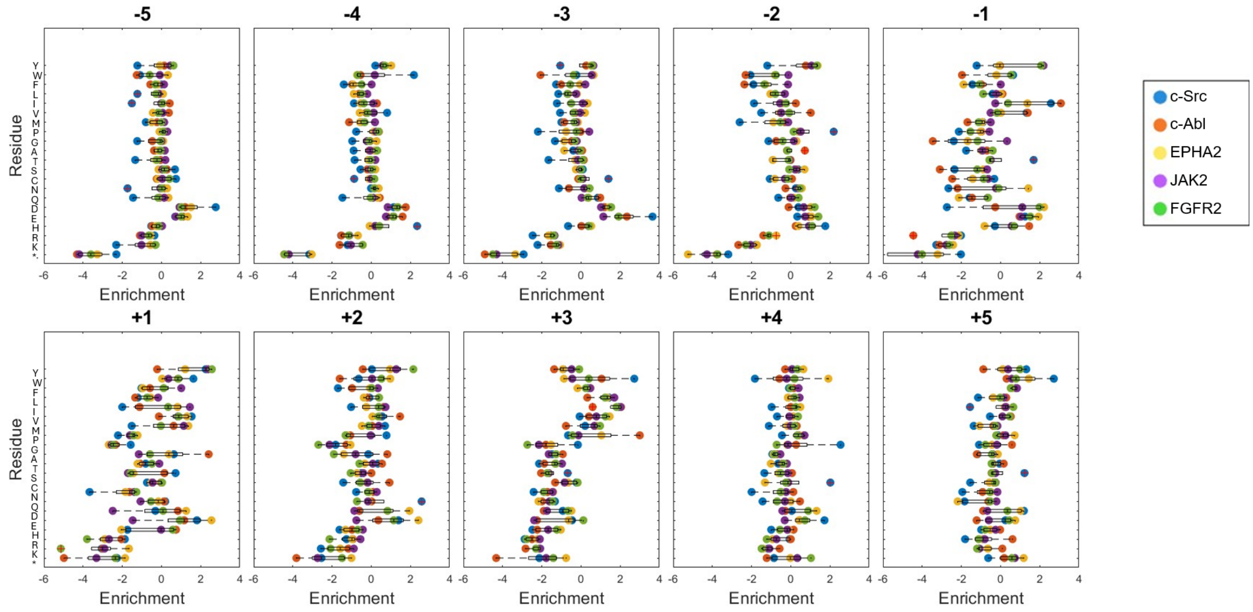


**Figure S13. Amino acid preferences for different sites.** The average enrichment of each amino acid at each substrate site ($X_5$-Y-$X_5$). Preferences differ most near the phospho-acceptor tyrosine (-2 through +3). All PTKs strongly prefer acidic residues on the N-terminus (-5 through -3).

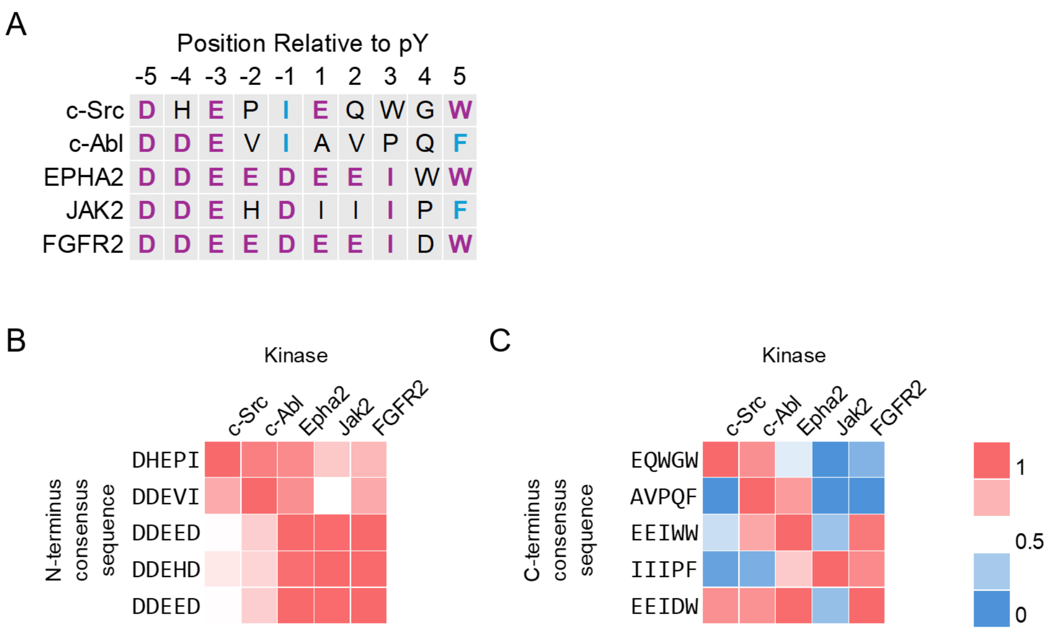


**Figure S14. Analysis of consensus peptides.** (A) Consensus peptides for each kinase. Shared residues appear in purple (most common) or blue (second most common). (B-C) Substrate compatibility ($P_{ij}$) scores for (B) the N-terminus and (C) the C-terminus show segments of the consensus peptides for each kinase. The C-terminus exhibits greater variability than the N-terminus.

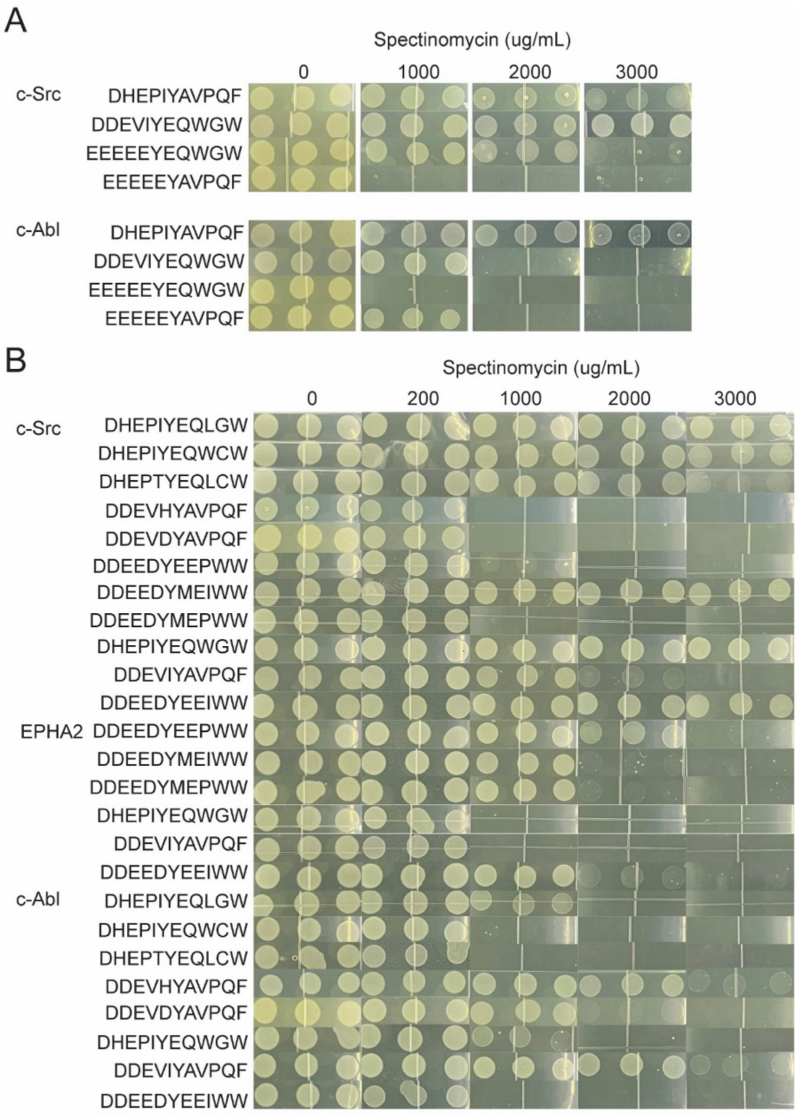


**Figure S15. Drop-based plating to examine PTK-substrate compatibility (Fig. 3).** Drop-based plating of s1030 cells harboring B2H systems (B2H$_{c\text{-}Src}$, B2H$_{c\text{-}Abl}$, and B2H$_{EPHA2}$) with the indicated substrates (pMM532 plasmids). Details: 3.5 uL drops of OD=0.1 cell culture placed on LB agar plates incubated at 37°C for 16-18 hours. Images: drops for n = 3 biological replicates.

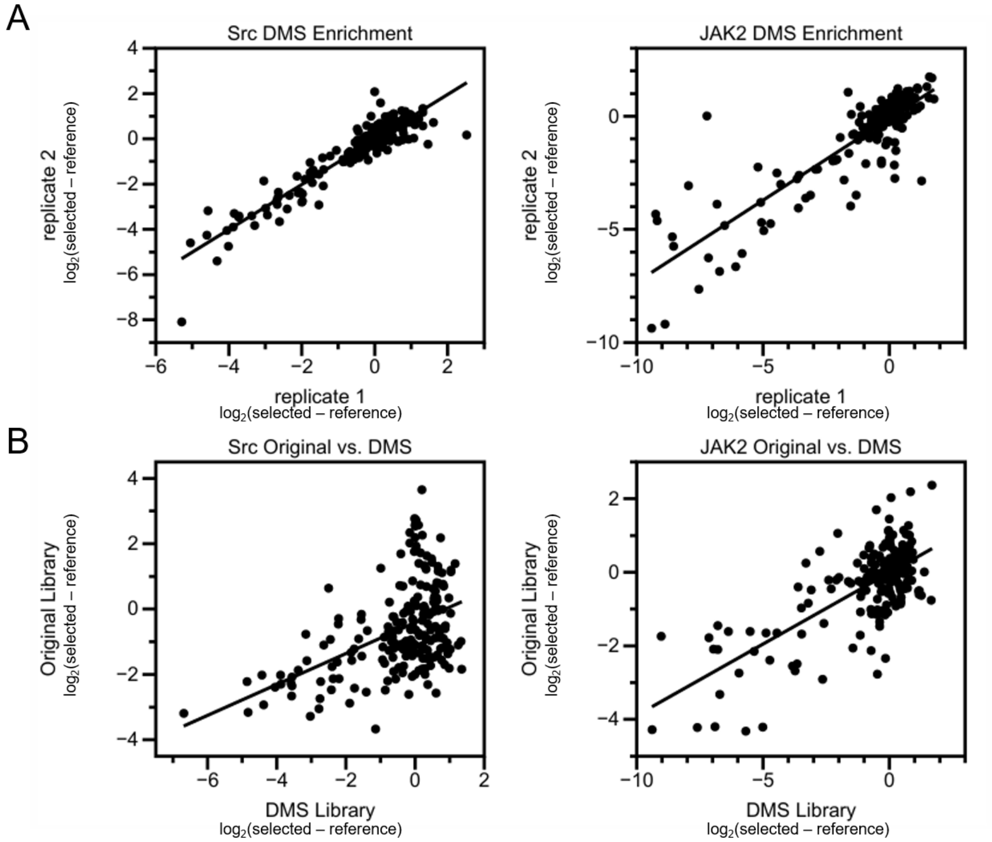


**Figure S16. Analysis of PSSMs based on DMS.** (A) These data show the reproducibility (i.e., $R^2$ for biological replicates) of 11-residue PSSMs built on deep mutational scanning (DMS) of the consensus substrates for c-Src and JAK2 (Fig. 4A). (B) Comparison of the average enrichment scores for each residue at each site in the random (Fig. 2A) and DMS (Fig. 4A) libraries. Discrepancies between the datasets suggest context-dependent preferences.

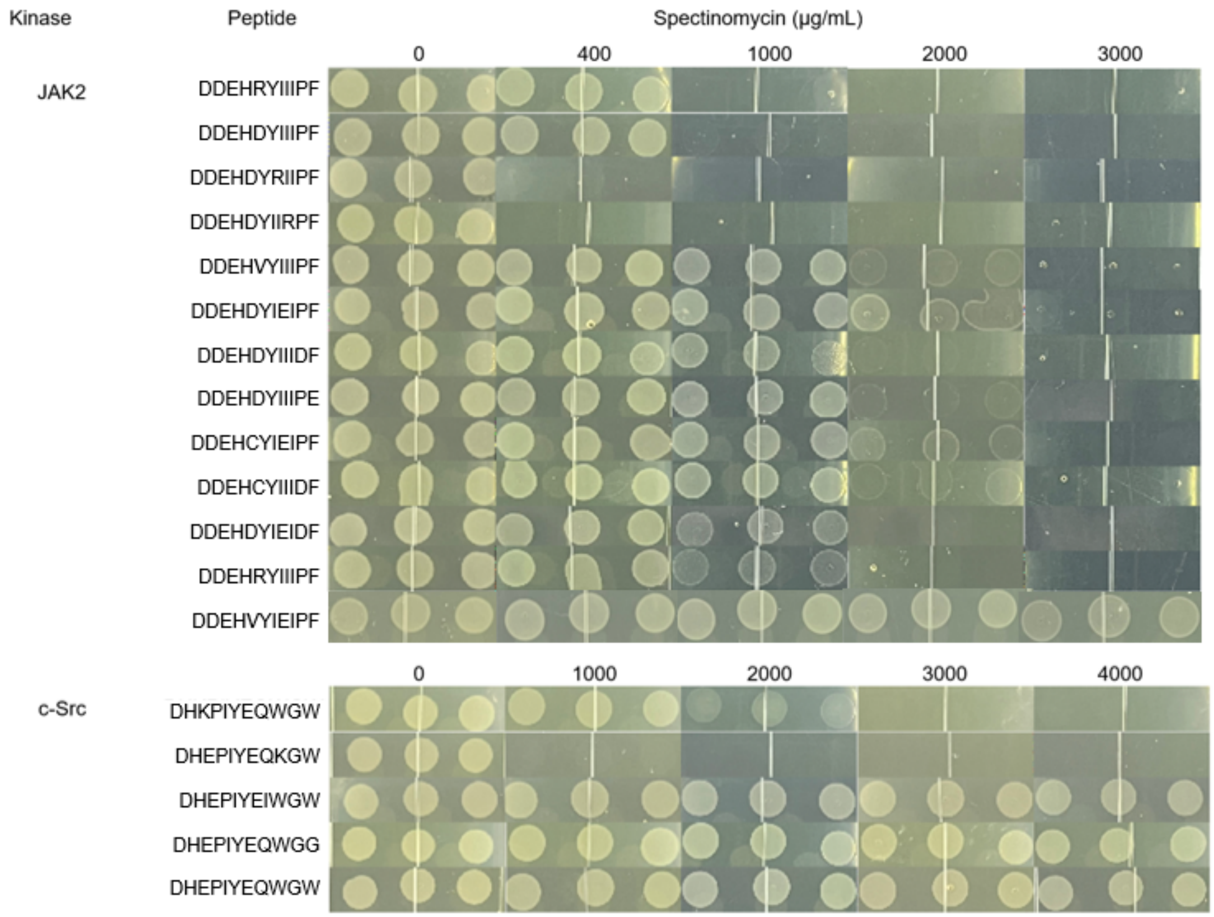


**Figure S17. Drop-based plating to examine PTK-substrate compatibility.** Drop-based plating of s1030 cells harboring B2H systems ($B2H_{JAK2}$ and $B2H_{cSrc}$) with the indicated substrates (pMM532 plasmids). Details: For c-Src, we used 3.5 uL drops of OD=0.1 cell culture placed on LB agar plates incubated at 37°C for 16-18 hours. For JAK2, we used 30°C and 24 hours. Images: drops for n = 3 biological replicates.

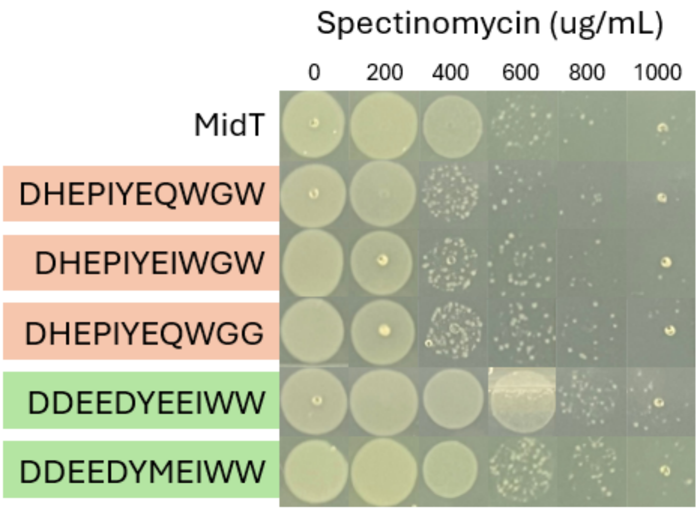


**Figure S18. Examining peptide mutants in a B2H system with reduced c-Src expression.** Deep mutational scanning (DMS) results suggest that +2Ile and +5Gly mutations to the c-Src consensus (pink) will further improve compatibility with c-Src (Fig. 4A) while the mutation +1Met on the EPHA2 consensus peptide (green) should decrease compatibility (Fig. 3D). In our original B2H system, c-Src confers survival to very high spectinomycin levels, making small changes in compatibility difficult to detect. Here, we reduced c-Src expression levels by swapping in the proD promoter for the prom46 promoter. In this more sensitive system, improvements in the compatibility of the c-Src consensus motif (pink), as measured by spectinomycin resistance, remain difficult to detect; however, a reduction in the compatibility with the +1Met variant of the EPHA2 sequence (green) is readily apparent.

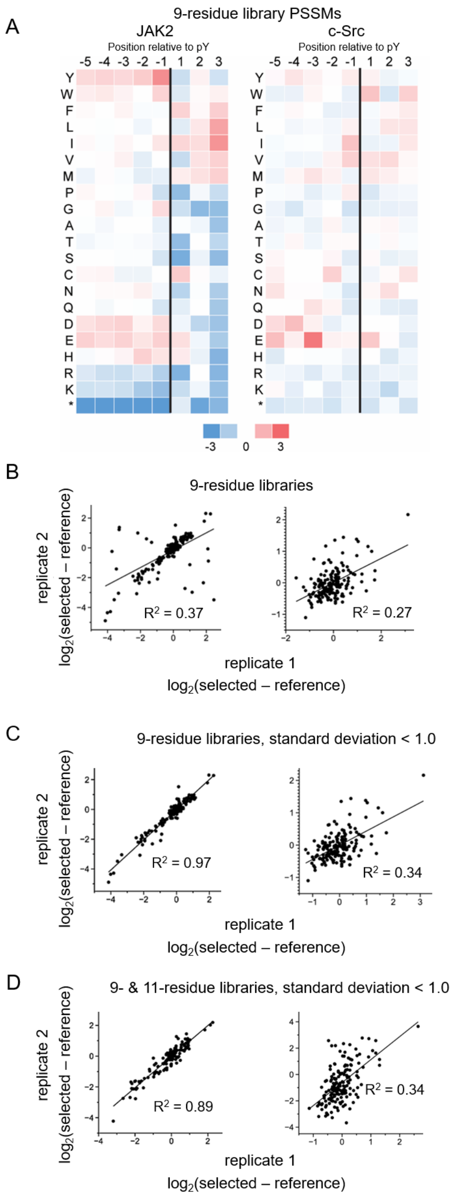


**Figure S19. PSSMs for 9-residue substrate libraries.** (A) We screened 9-residue peptide libraries against JAK2, which showed stop codon enrichment at the +4 and +5 poistions of 11-residue substrates, and c-Src, which did not. Numbers show the average enrichment of each amino acid at each position for n = 2 biological replicates. (B-C) Comparisons of biological replicates for (B) the 9-residue library and (C) the 9-residue library showing only highly reproducible datapoints (SD < 1). Left: JAK2. Right: c-Src. (D) Comparison of highly reproducible enrichment scores for 9- and 11-residue libraries for each PTK.

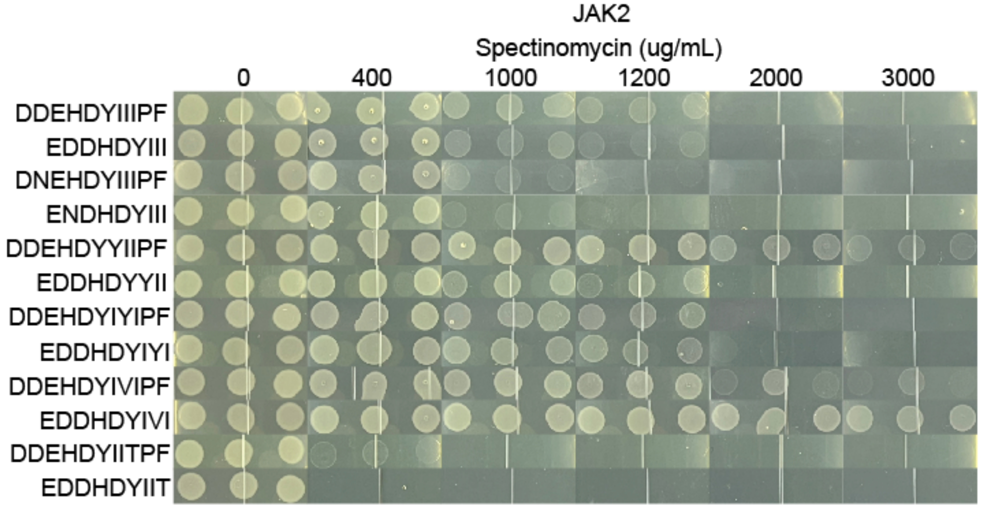


**Figure S20. Drop-based plating to examine PTK-substrate compatibility (Fig. 4B).** Drop-based plating of s1030 cells harboring B2H systems ($B2H_{JAK2}$) with the indicated substrates (pMM532 plasmids). Details: 3.5 uL drops of OD=0.1 cell culture placed on LB agar plates incubated at 30°C for 24 hours. Images: drops for n = 3 biological replicates.

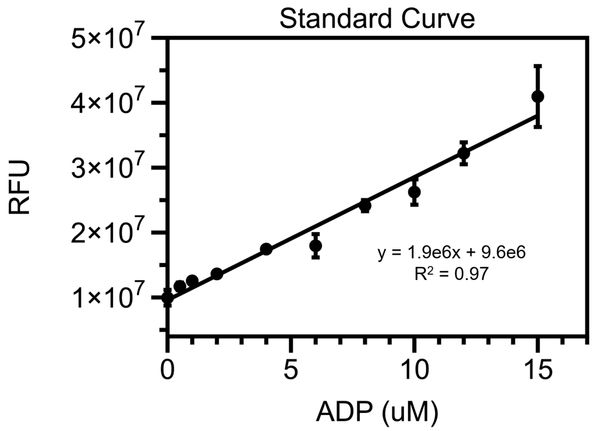


**Figure S21. Standard curve for ADP formation.** Using the ADP Quest assay (Eurofins, see Methods), we measured fluorescence ($\lambda_{ex}$ = 530 / $\lambda_{em}$ = 590) at set ADP concentrations to generate a standard curve. Data depict the mean and standard deviation for n = 5 measurements.

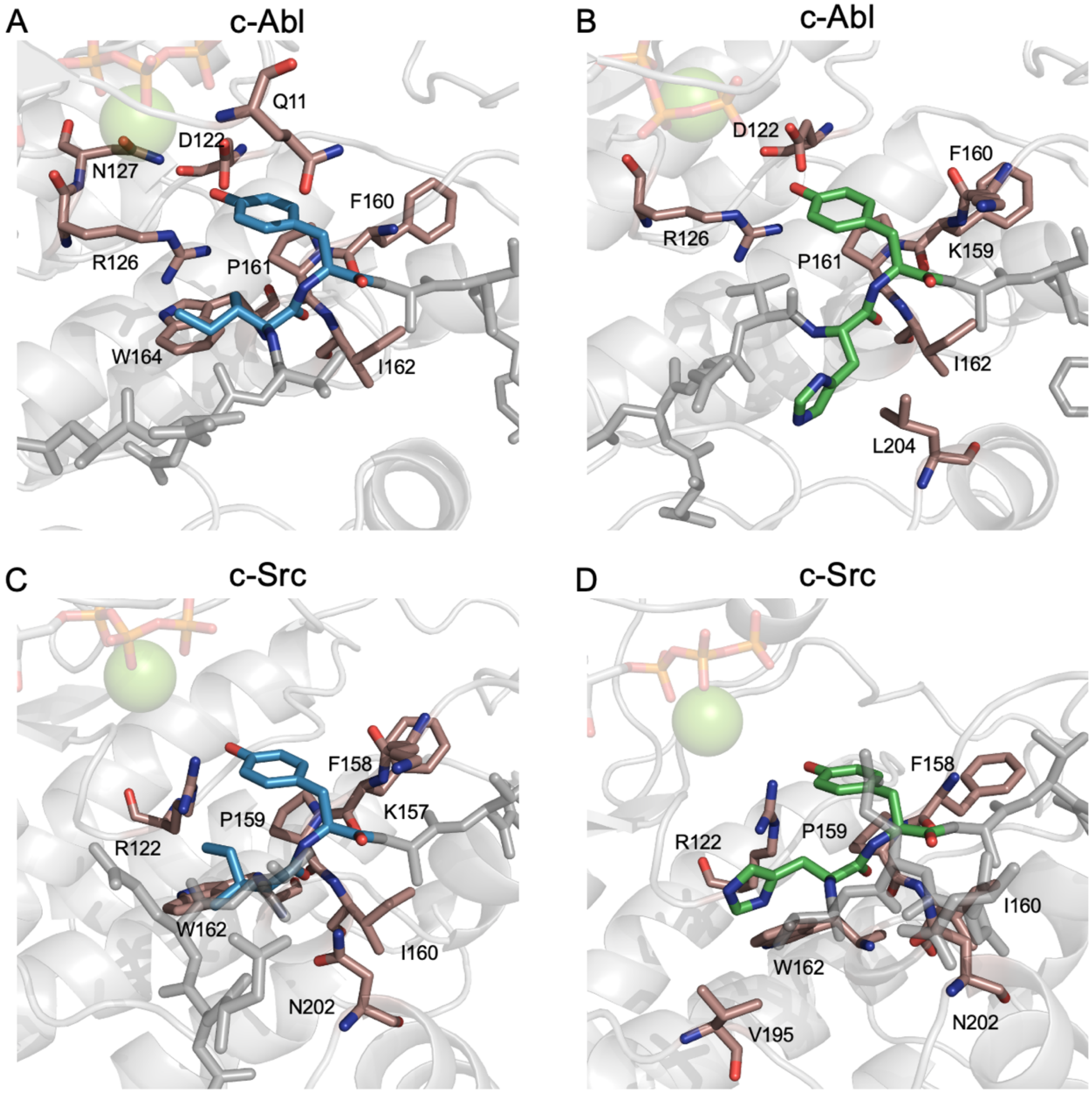


**Figure S22. AlphaFold 3.0 (AF3) of PTK-substrate compatibility (Fig. 5).** (A-B) AF3 structures of c-Abl bound to (A) the c-Abl consensus (DDEVIYAVPQF, blue) and (B) its -1His mutant (DDEVHYAVPQF, green). (C-D) AF3 structures of c-Src bound to (C) the c-Abl consensus and (D) its -1His mutant.

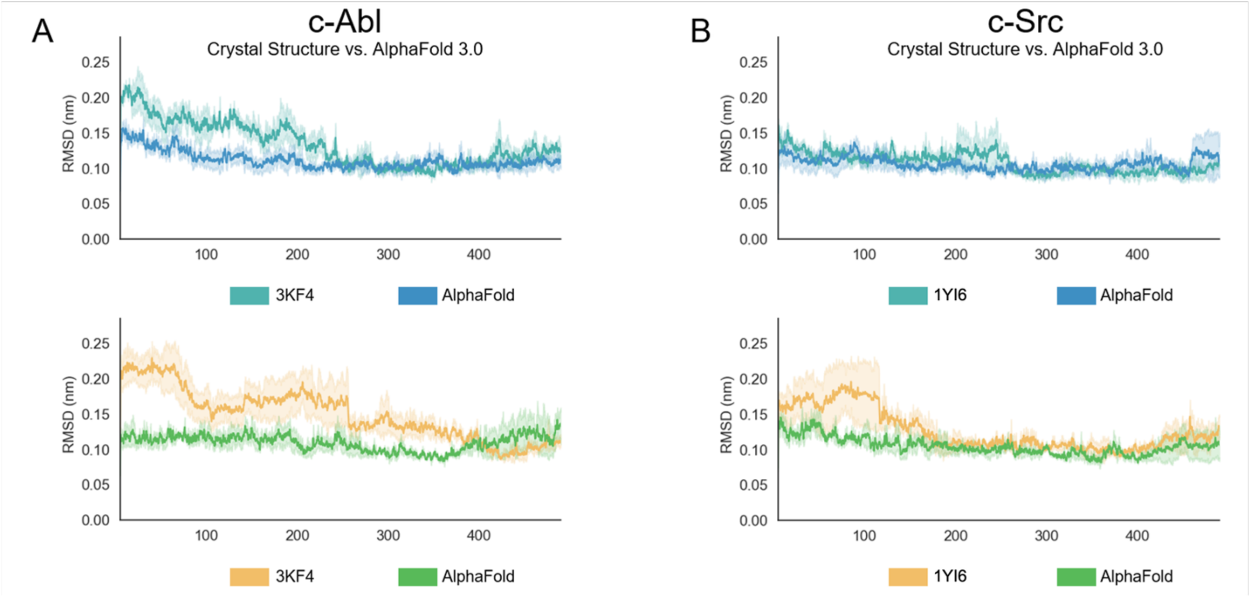


**Figure S23. Root-mean-square deviation of PTK backbone atoms from their respective centroids.** (A) RMSD values for c-Abl and the c-Abl consensus peptide (DDEVIYAVPQF; top) and -1His mutant (DDEVHYAVPQF, bottom). (B) RMSD values for c-Src and the c-Abl consensus peptide (top) and -1His mutant (bottom). All plots show runs seeded from crystal structures (PDB entries) or top-scoring AF3 predictions (AlphaFold). For c-Src, these preliminary simulations used a glutamate substitution for the pTyr at position 416, but final simulations with AF3-seeded structures and the pTyr at that position showed no significant deviations from the preliminary data. For both PTKs, runs seeded by AlphaFold predictions exhibited similar or lower RMSDs. Data depict averages of n=3 replicates with standard error.

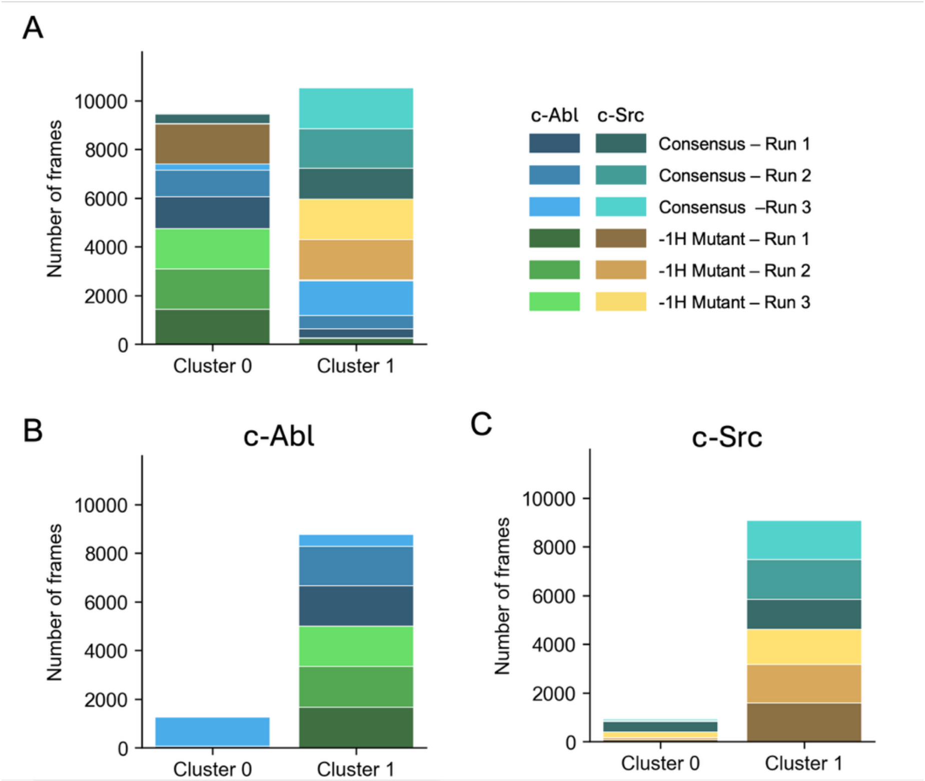


**Figure S24. Clustering of peptide backbone structures for both inter- and intra- PTK trajectory frames.** (A) K-means clustering (n = 3) of all MD trajectories of c-Abl and c-Src bound to the c-Abl consensus and -1His mutant peptides with individual replicates resolved (versus all replicates combined, as in Fig. 5F). Clusters separated c-Abl peptides from c-Src peptides but not compatible peptides from incompatible peptides. (B-C) Clustering carried out separately for (B) c-Abl trajectories and (C) c-Src trajectories. For both PTKs, clustering did not distinguish between the two peptide structures or group them by PTK compatibility; the largest cluster contained similar numbers of frames from both.

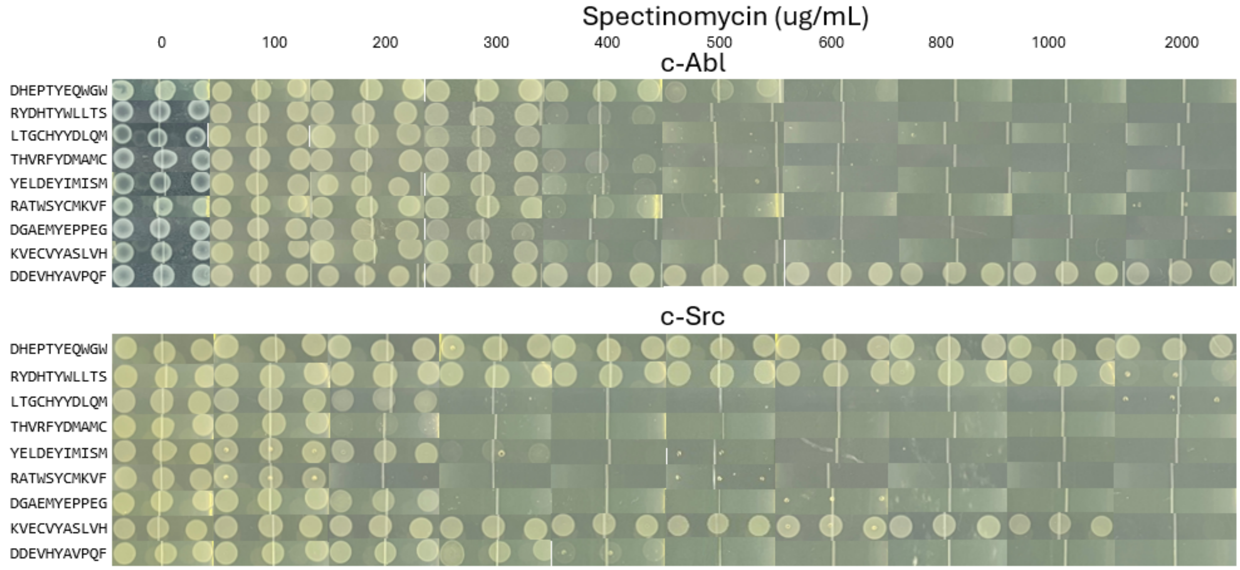


**Figure S25. Drop-based plating to examine PTK-substrate compatibility (Fig. 6B).** Drop-based plating of s1030 cells harboring B2H systems ($B2H_{cSrc}$ and $B2H_{cSrc}$) with the indicated substrates (pMM532 plasmids). Details: We used 3.5 uL drops of OD=0.1 cell culture placed on LB agar plates incubated at 37°C for 16-18 hours.

### SI Tables

#### Table S1. Human PTKs grouped by substrate specificity via hierarchical clustering

| *PTK* | *Phylogeny** [20]* | *Specificity Group*** [18]* |
|---|---|---|
| *c-Src* | Src module | Src |
| *c-Abl* | Src module | Abl |
| *EPHA2* | Eph | Ephrin receptors |
| *JAK2* | Jak | JAK |
| *FGFR2* | Growth factor receptors | FGF/VEGF receptors |

**Phylogeny* lists groups identified by Wayland et al. [20]

***Specificity Group* refers to the group determined by using hierarchical clustering to group human PTKs by substrate specificity in Yaron-Barir et al. [18]

**Table S2. A comparison of compatibility scores and growth-based plating.** Data show the peptide scores, as described by Li et al. [6] (SI note 1) using our B2H-based PSSMs, and the maximum antibiotic resistance conferred by each PTK-substrate pair in our growth experiments.

| Substrate | Score | ug/mL |
|---|---|---|
| DHEPIYEQWGW | 1.00 | 3000 |
| DHEPIYEQWCW | 0.98 | 4000 |
| DHEPTYEQWGW | 0.97 | 3000 |
| DHEPIYEQLGW | 0.96 | 3000 |
| DHEPIYEIWGW | 0.89 | 4000 |
| DHEPIYEQWGG | 0.88 | 4000 |
| DDEVIYEQWGW | 0.86 | 3000 |
| DHEPIYEQKGW | 0.85 | 0 |
| DHKPIYEQWGW | 0.82 | 2000 |
| EEEEEYEQWGW | 0.80 | 3000 |
| DDEEDYEEIDW | 0.63 | 3000 |
| DHEPIYAVPQF | 0.62 | 3000 |
| DDEEDYEEIWW | 0.55 | 3000 |
| DDEEDYSEIWW | 0.52 | 3000 |
| DDEEDYEDIWW | 0.49 | 0 |
| DDEEDYEEVWW | 0.49 | 3000 |
| DDEVIYAVPQF | 0.48 | 3000 |
| DDEEDYEEPWW | 0.48 | 1000 |
| RYDHTYWLLTS | 0.46 | 2000 |
| DDEEDYMEIWW | 0.45 | 3000 |
| DDEEDYIEIWW | 0.44 | 2000 |
| EEEEEYAVPQF | 0.42 | 0 |
| DDEVHYAVPQF | 0.40 | 400 |
| DDEHDYIIIPF | 0.39 | 0 |
| DDEVDYAVPQF | 0.32 | 0 |
| DGAEMYEPPEG | 0.32 | 400 |
| KVECVYASLVH | 0.28 | 1000 |
| YELDEYIMISM | 0.16 | 300 |
| LTGCHYYDLQM | 0.08 | 200 |
| THVRFYDMAMC | 0.02 | 200 |
| RATWSYCMKVF | 0.00 | 100 |

| Substrate | Score | ug/mL |
|---|---|---|
| DDEVIYAVPQF | 1.00 | 3000 |
| DDEVHYAVPQF | 0.94 | 3000 |
| DHEPIYAVPQF | 0.92 | 3000 |
| EEEEEYAVPQF | 0.91 | 1000 |
| DDEVDYAVPQF | 0.88 | 2000 |
| DDEVIYEQWGW | 0.80 | 1000 |
| DDEEDYEEPWW | 0.73 | NA |
| DHEPIYEQWGW | 0.72 | 1000 |
| DHEPIYEQWCW | 0.71 | 0 |
| EEEEEYEQWGW | 0.71 | 0 |
| DHEPIYEQLGW | 0.69 | 1000 |
| DDEEDYEEIDW | 0.65 | 0 |
| DDEEDYEEIWW | 0.64 | NA |
| DDEEDYEEVWW | 0.63 | NA |
| DDEEDYWEIWW | 0.62 | NA |
| DDEEDYMEIWW | 0.62 | NA |
| DDEEDYEDIWW | 0.60 | NA |
| DDEEDYSEIWW | 0.60 | NA |
| DDEHDYIIIPF | 0.59 | 0 |
| DHEPTYEQWGW | 0.59 | 500 |
| KVECVYASLVH | 0.57 | 500 |
| DDEEDYIEIWW | 0.55 | NA |
| DGAEMYEPPEG | 0.52 | 400 |
| YELDEYIMISM | 0.39 | 500 |
| RYDHTYWLLTS | 0.36 | 400 |
| LTGCHYYDLQM | 0.36 | 300 |
| THVRFYDMAMC | 0.25 | 500 |
| RATWSYCMKVF | 0.04 | 500 |
| MSTKAYKRAQR | 0.03 | TBD |
| PPKSRYNPTN* | 0.00 | TBD |

| Substrate | Score | ug/mL |
|---|---|---|
| DDEEDYEEIWW | 1.00 | 2000 |
| DDEEDYEDIWW | 0.96 | 3000 |
| DDEEDYEEVWW | 0.95 | 2000 |
| DDEEDYEEIDW | 0.95 | 2000 |
| DDEEDYEEPWW | 0.92 | 2000 |
| DDEEDYMEIWW | 0.91 | 1000 |
| DDEEDYWEIWW | 0.80 | 1000 |
| DDEEDYIEIWW | 0.69 | 1000 |
| DDEEDYSEIWW | 0.48 | 0 |
| DDEHDYIIIPF | 0.44 | 1000 |
| DHEPIYEQLGW | 0.36 | NA |
| EEEEEYEQWGW | 0.33 | NA |
| EEEEEYAVPQF | 0.29 | NA |
| DDEVDYAVPQF | 0.24 | NA |
| DDEVIYEQWGW | 0.23 | NA |
| DHEPIYEQWGW | 0.19 | 0 |
| DDEVIYAVPQF | 0.17 | 0 |
| DHEPIYAVPQF | 0.13 | NA |
| DHEPIYEQWCW | 0.10 | NA |
| DHEPTYEQWGW | 0.04 | NA |
| DDEVHYAVPQF | 0.00 | NA |

| Substrate | Score | ug/mL |
|---|---|---|
| EDDHDYYII | 1.00 | 1000 |
| EDDHDYIYI | 0.92 | 2000 |
| DDEHDYYIIPF | 0.90 | 3000 |
| DDEHDYIYIPF | 0.83 | 1000 |
| EDDHDYIII | 0.80 | 1000 |
| DDEHDYIIIPF | 0.73 | 1000 |
| EDDHDYIVI | 0.72 | 3000 |
| DDEHDYIVIPF | 0.67 | 3000 |
| ENDHDYIII | 0.64 | 1000 |
| DNEHDYIIIPF | 0.61 | 1000 |
| DDEHDYIIIDF | 0.53 | 2000 |
| DDEHDYIIIPE | 0.48 | 2000 |
| DDEHDYIEIPF | 0.47 | 3000 |
| DDEHVYIIIPF | 0.45 | 2000 |
| DDEHDYIIIPR | 0.44 | 400 |
| DDEHDYIITPF | 0.39 | 400 |
| DDEEDYEEIWW | 0.36 | 0 |
| DDEHDYIEIDF | 0.27 | 1000 |
| DDEHCYIIIDF | 0.25 | 2000 |
| DDEHCYIEIPF | 0.18 | 2000 |
| EDDHDYIIT | 0.13 | 0 |
| DDEHRYIIIPF | 0.12 | 400 |
| DDEVIYAVPQF | 0.08 | 0 |
| DDEHDYRIIPF | 0.00 | 0 |
| DDEHDYIIRPF | 0.00 | 0 |
| DHEPIYEQWGW | 0.00 | 0 |

### Table S3. Gene Sources.

| *Component* | *Organism* | *Plasmid* | *Source* |
|---|---|---|---|
| *c-Src* | *H. sapiens* | pDONR223_SRC_WT | AG: 82165 |
| *c-Abl* | *M. musculus* | N/A | Residues 223-496 codon optimized for expression in *E. coli* |
| *EPHA2* | *H. sapiens* | N/A | Residues 590-976 codon optimized for expression in *E. coli* |
| *JAK2* | *H. sapiens* | N/A | Residues 835:1132 codon optimized for expression in *E. coli* |
| *FGFR2* | *H. sapiens* | N/A | Residues 365:768 codon optimized for expression in *E. coli* |
| *B2H system parts (cI_SH2, CDC37, SpecR, RpoZ)* | *N/A* | pB2H | Sarkar et al.[4] |
| *sfGFP* | *N/A* | pBAD pduP1-18-gfpmut2-ssrA | AG: 69512 |

### Table S4. Plasmids used in this study.

| *Plasmid* | *Description* | *Antibiotic** | *Availability* |
|---|---|---|---|
| *F-plasmid* | The F-plasmid from the S1030 strain of *E. coli*. | T | AG: 105063[‡] |
| *$B2H_{c\text{-}Src}$* | Bacterial two-hybrid system. Contains cI_SH2, c-Src, CDC37, SpecR. | K, S | Fox Lab |
| *$B2H_{c\text{-}Src,\ ProD}$* | Bacterial two-hybrid system. Contains cI_SH2, c-Src under proD promoter, CDC37, SpecR | K,S | Fox Lab |
| *$B2H_{c\text{-}Abl}$* | Bacterial two-hybrid system. Contains cI_SH2, c-Abl, CDC37, SpecR | K, S | Fox Lab |
| *$B2H_{EPHA2}$* | Bacterial two-hybrid system. Contains cI_SH2, EPHA2, CDC37, SpecR | K,S | Fox Lab |
| *$B2H_{JAK2}$* | Bacterial two-hybrid system. Contains cI_SH2, JAK2, CDC37, SpecR | K,S | Fox Lab |
| *$B2H_{FGFR2}$* | Bacterial two-hybrid system. Contains cI_SH2, FGFR2, CDC37, SpecR | K,S | Fox Lab |
| *pMM532* | Plasmid constitutively expressing rpoZ-substrate in place of an HrtR gene under a ProD promoter. | C | AG: 112529 |
| *$pMM532_{S}$* | pMM532 constitutively expressing rpoZ-MidT in place of HrtR gene. | C | Fox Lab |
| *$pMM532_{S\_Y/F}$* | pMM532 constitutively expressing MidT Y/F in place of HrtR gene. | C | Fox Lab |

| *Plasmid* | *Description* | *Antibiotic** | *Availability* |
|---|---|---|---|
| *pMM532*$_{Substrate\_Library}$ | pMM532 constitutively expressing rpoZ-S where S is substrate library $X_5$-Y-$X_5$ with 290,000 variants. | C | Fox Lab |
| *pMM532*$_{9\text{-}residue\_Library}$ | pMM532 constitutively expressing rpoZ-S where S is substrate library $X_5$-Y-$X_3$ with 700,000 variants | C | Fox Lab |
| *pMM532*$_{Src\_consensus\_DMS\_Library}$ | pMM532 cloned to contain DMS library randomizing individual residues from c-Src consensus DHEPIYEQWGW | C | Fox Lab |
| *pMM532*$_{JAK2\_consensus\_Library}$ | pMM532 cloned to contain DMS library randomizing individual residues from JAK2 consensus DDEHDYIIIPF | C | Fox Lab |
| *pMM532*$_{c\text{-}Src\_consensus}$ | pMM532 constitutively expression rpoZ-c-Src consensus peptide DHEPIYEQWGW in place of HrtR gene. | C | Fox Lab |
| *pMM532*$_{c\text{-}Abl\_consensus}$ | pMM532 constitutively expression rpoZ-c-Abl consensus peptide DDEVIYAVPQF in place of HrtR gene. | C | Fox Lab |
| *pMM532*$_{EPHA2\_consensus}$ | pMM532 constitutively expression rpoZ-EPHA2 consensus peptide DDEEDYEEIWW in place of HrtR gene. | C | Fox Lab |
| *pMM532*$_{JAK2\_consensus}$ | pMM532 constitutively expression rpoZ-JAK2 consensus peptide DDEHDYIIIPF in place of HrtR gene. | C | Fox Lab |
| *pMM532*$_{FGFR2\_consensus}$ | pMM532 constitutively expression rpoZ-FGFR2 consensus peptide DDEEDYEEIDW in place of HrtR gene. | C | Fox Lab |

*Antibiotic resistance: carbenicillin (C, 50 μg/ml), kanamycin (K, 50 μg/ml), tetracycline (T, 10 μg/ml), chloramphenicol (P, 34 μg/ml), and spectinomycin (S, conditional).
‡AG=Addgene accession # (Addgene.com).

### Table S5. Primers used for plasmid assembly and NGS sequencing.

| *Component* | *F Primer* | *R Primer* |
|---|---|---|
| *Substrate_Library for restriction digest into pMM532 plasmid* | TTCTAGAGCACAGCTAACACCAC | ATATCTCGAGTTAMNNMNNMNNMNNMNNATAMNNMNNMNNMNNMNNCGCAGCTGCACGAC |
| *Backbone for pMM532 for individual substrate insertions* | CGTAAACTTGGTCTGACAGTTACC | CGCAGCTGCACGACGACC |
| *Insert for DWEHIYEEICW substrate into pMM532* | GATTGGGAGCATATTTATGAGGAGATTTGTTGGTAATAACTCGAGGGATCCCATGGT | GGTAACTGTCAGACCAAGTTTACG |

| Component | F Primer | R Primer |
|---|---|---|
| Insert for CDCPEYWQLEY substrate into pMM532 | TGTGATTGTCCGGAGTATTGGCAGCTGGAGTATTAATAACTCGAGGGATCCCATGGT | GGTAACTGTCAGACCAAGTTTACG |
| Insert for DDEVHYAVPQF substrate into pMM532 | GATGATGAGGTGCATTATGCAGTGCCGCAGTTTTAATAACTCGAGGGATCCCATGGT | GGTAACTGTCAGACCAAGTTTACG |
| Insert for EPHA2_CONSENSUS into pMM532 | GATGATCATGAGGAGTATTATGATCCGTGGTGGTAATAACTCGAGGGATCCCATGGT | CCACCACGGATCATAATACTCCTCATGATCATCCGCAGCTGCACGACGACC |
| Insert for JAK2_CONSENSUS into pMM532 | GGTCGTCGTGCAGCTGCGGATGATGAGCATGATTATATTATTATTCCGTTTTAATAACTCGAGGGATCCCATGGT | GGTAACTGTCAGACCAAGTTTACG |
| Insert for FGFR2_CONSENSUS into pMM532 | GGTCGTCGTGCAGCTGCGGACGACGAGGAGGACTATGAGGAGATTGACTGGTAATAACTCGAGGGATCCCATGGT | GGTAACTGTCAGACCAAGTTTACG |
| Insert for XXXXXYXXX into pMM532 | GGTCGTCGTGCAGCTGCGMNNMNNMNNMNNMNNTATMNNMNNMNNTAATAACTCGAGGGATCCCATGGT | GGTAACTGTCAGACCAAGTTTACG |
| NGS sequencing amplicon | TCCCTACACGACGCTCTTCCGATCTCGGAAGAAAACGATAAAACCACTG | GTTCAGACGTGTGCTCTTCCGATCTCACGCGTACCATGGGATCC |

*Individual mutations added to peptide sequences above, keeping melting temperatures and binding sequences consistent

### Table S6. Library sizes

| Library | Estimated plated library size |
|---|---|
| $X_5$-Y-$X_5$ | 290,000 |
| $X_5$-Y-$X_3$ | 700,000 |
| DHEPIYEQWGW DMS | 4,500 |
| DDEHDYIIIPF DMS | 5,000 |
| c-Src$^+$, c-Abl$^-$ | 100 |

**Table S7.** Consensus peptides.

| PTK | Sequence |
|---|---|
| c-Src | DHEPIYEQWGW |
| c-Abl | DDEVIYAVPQF |
| EPHA2 | DDEEDYEEIWW |
| JAK2 | DDEHDYIIIPF |
| FGFR2 | DDEEDYEEIDW |

**Table S8. Enrichment scores for heat maps.** The accompanying Excel file provides enrichment scores for all PSSMs, obtained by screening $pMM532_{substrate\_library}$, $pMM532_{9\text{-}residue\_library}$, $pMM532_{c\text{-}Src\_consensus\_library}$, or $pMM532_{JAK2\_consensus\_library}$ against s1030 cells + $B2H_{c\text{-}Src}$, $B2H_{c\text{-}Abl}$, $B2H_{EPHA2}$, $B2H_{JAK2}$, or $B2H_{FGFR2}$. Libraries plated on LB agar plates and grown 37°C for 16-18 hours (c-Src, c-Abl, EPHA2, and FGFR2) or 30°C for 48 hours (JAK2).

**Table S9. Cosine similarity scores.** The accompanying Excel file contains all cosine similarity scores in all main and supplementary figures as described in the Methods section and Figure S7.

**Table S10. Kinetics data.** The accompanying Excel file provides measurements for all kinetics measurements made in this study, including standard error and sample sizes.
